# Systemic trafficking of mRNA lipid nanoparticle vaccine following intramuscular injection generates potent tissue-specific T cell response

**DOI:** 10.1101/2025.04.21.649878

**Authors:** Christine Wei, Yining Zhu, Xiaoya Lu, Kailei D. Goodier, Di Yu, Xiang Liu, Joseph Choy, Alejandro Téllez Calderón, Jingyao Ma, Yunhe Su, Jinghan Lin, Shuyi Li, Jonathan P. Schneck, Sean C. Murphy, Hai-Quan Mao

**Affiliations:** Department of Biomedical Engineering, Johns Hopkins University School of Medicine, Baltimore, MD 21218, USA; Institute for NanoBioTechnology, Johns Hopkins University, Baltimore, MD 21218, USA; Translational Tissue Engineering Center, Johns Hopkins University School of Medicine, Baltimore, MD 21218, USA; Department of Materials Science and Engineering, Johns Hopkins University, Baltimore, MD 21218, USA; Department of Chemical and Biomolecular Engineering, Johns Hopkins University, Baltimore, MD 21218, USA; Institute for Cell Engineering, Johns Hopkins University School of Medicine, Baltimore, MD 21218, USA; Department of Pathology, Johns Hopkins University School of Medicine, Baltimore, MD 21218, USA; Department of Medicine, Johns Hopkins University School of Medicine, Baltimore, MD 21218, USA; Department of Oncology, Johns Hopkins University School of Medicine, Baltimore, MD 21218, USA; Department of Laboratory Medicine and Pathology, University of Washington, Seattle, WA 98109, USA; Department of Emerging and Re-emerging Infectious Diseases, University of Washington, Seattle, WA 98109, USA; Department of Microbiology, University of Washington, Seattle, WA 98109, USA

**Author notes:** Corresponding authors’,. These authors contributed equally to this work (Christine Wei, Yining Zhu).

## Abstract

The mRNA lipid nanoparticles (LNPs) represent a new generation of vaccine carriers designed to elicit potent immune responses against infectious diseases and cancer. Despite the clinical success and rapid advancements in mRNA LNP technologies, the trafficking patterns of LNPs after intramuscular (i.m.) administration and the subsequent tissue-specific immunological effects have not been systematically characterized. Here, we report that trafficking of mRNA LNPs to different organs following i.m. injection is crucial for the induction of tissue-specific immunity beyond systemic immune response, particularly in tissue-resident CD8^+^ T cell generation, which is important for localized defense. By fine-tuning the composition of mRNA LNPs, trafficking patterns to systemic organs can be modulated, which can alter the resulting tissue-specific immune response. Formulations with a greater ability to enter the bloodstream can preferentially localize and transfect cells in specific organs like the liver, elicit stronger tissue-specific CD8^+^ T cell immune responses, and achieve enhanced efficacy in a liver tumor model. These findings highlight the potential to tailor mRNA LNP compositions to modulate trafficking following i.m. injection, thereby providing novel strategies for designing tissue-specific vaccines. Such strategies are particularly valuable for organ-specific diseases like cancer and infectious diseases, where tissue targeting and long-lasting immunity are essential for therapeutic success.

## Main

In infectious disease control and cancer therapy, there is increasing interest in achieving localized tissue-specific immunity through the induction of antigen-specific CD8^+^ T cells within the local tissue environment, such as CD8^+^ tissue-resident memory T (T_RM_) cells^1–4^. These tissue-specific T cells play a critical role in long-term immune surveillance and provide rapid responses upon pathogen re-encounter^5–8^. Their unique ability to remain localized and maintain a heightened state of readiness makes them particularly effective at controlling pathogens in specific tissues^9–12^. Consequently, the induction of such tissue-specific CD8^+^ T cells has become a key consideration in the development of vaccines targeting tissue-specific infections^13^.

Antigen-expression within the tissue microenvironment facilitates recruitment of antigen-specific CD8^+^ T cells to the tissue. Upon antigen recognition and adhesion receptor expression, a subset of these CD8^+^ T cells reside in the tissue in a persistent manner, maturing into the more specialized T_RM_ cell population^14,15^. This subset of T cells is optimized for localized immune surveillance, exhibiting minimal recirculation and can effectively control tissue-specific infection or tumor, offering an appealing strategy for long-term localized immunity, particularly for diseases where preventing infection at the organ level is crucial^1,16–18^. However, current antigen delivery strategies for achieving local tissue expression often rely on intravenous (i.v.) injection and complex targeting moieties, thus complicating vaccine development^1,19–22^. Furthermore, non-lymphoid tissue-specific transfection and the subsequent localized immune responses beyond systemic immunity via conventional vaccination methods, such as intramuscular (i.m.) injections, remains underexplored.

Lipid nanoparticles (LNPs) now serve as widely used COVID-19 vaccine delivery platforms for antigen-encoding mRNA and hold promise for additional prophylactic and therapeutic applications^23–25^. Recent studies demonstrated that the composition of mRNA LNP formulations impacts their pharmacokinetics^26–30^. The biodistribution and transfection behavior of LNPs administered by various routes—such as i.m., i.v., and subcutaneous (s.c.) injections—can be engineered by modifying their composition, thereby achieving tissue- and cell-specific transgene expression^22,26–34^. Additionally, adjusting LNP composition can influence the profile of transfected cell types, altering immune activation patterns, which ultimately impact vaccination outcomes^34–36^. On the other hand, an increasing number of recent studies have reported that a portion of certain LNPs administered via i.m. injection enters the systemic circulation and accumulates in major organs like the liver and lungs^37–40^. However, the therapeutic implications of LNP trafficking and unintended expression in non-lymphoid organs remain poorly understood.

Here, we hypothesized that tissue-specific CD8^+^ T cell immune responses could be generated by optimizing LNP compositions to promote preferential entry and expression in different non-lymphoid organs, including the liver and lungs (**Fig. 1a**). We first investigated the trafficking of mRNA LNPs following i.m. injection, focusing on their systemic entry and retention in non-lymphoid tissues, as well as the resulting systemic and tissue-specific immunological outcomes. We then examined the pathways underlying tissue-specific immune responses generated by i.m. injection of mRNA LNPs and their relationships with LNP trafficking and tissue-specific antigen expression. By further assessing the biodistribution profiles of different mRNA LNP compositions, we identified formulations with a higher degree of expression in either the liver or lungs, demonstrating the ability to modulate LNP trafficking and local expression by adjusting LNP compositions. These formulations were further evaluated for their short- and long-term immune profiles, including systemic cellular and humoral immune responses, as well as tissue-specific CD8^+^ T cell activation and CD8^+^ T_RM_ cell generation. Finally, we revealed the feasibility of tuning mRNA LNP formulation to induce durable systemic and localized immunity with improved therapeutic and prophylactic outcomes using mouse liver cancer models.

**Fig. 1:**
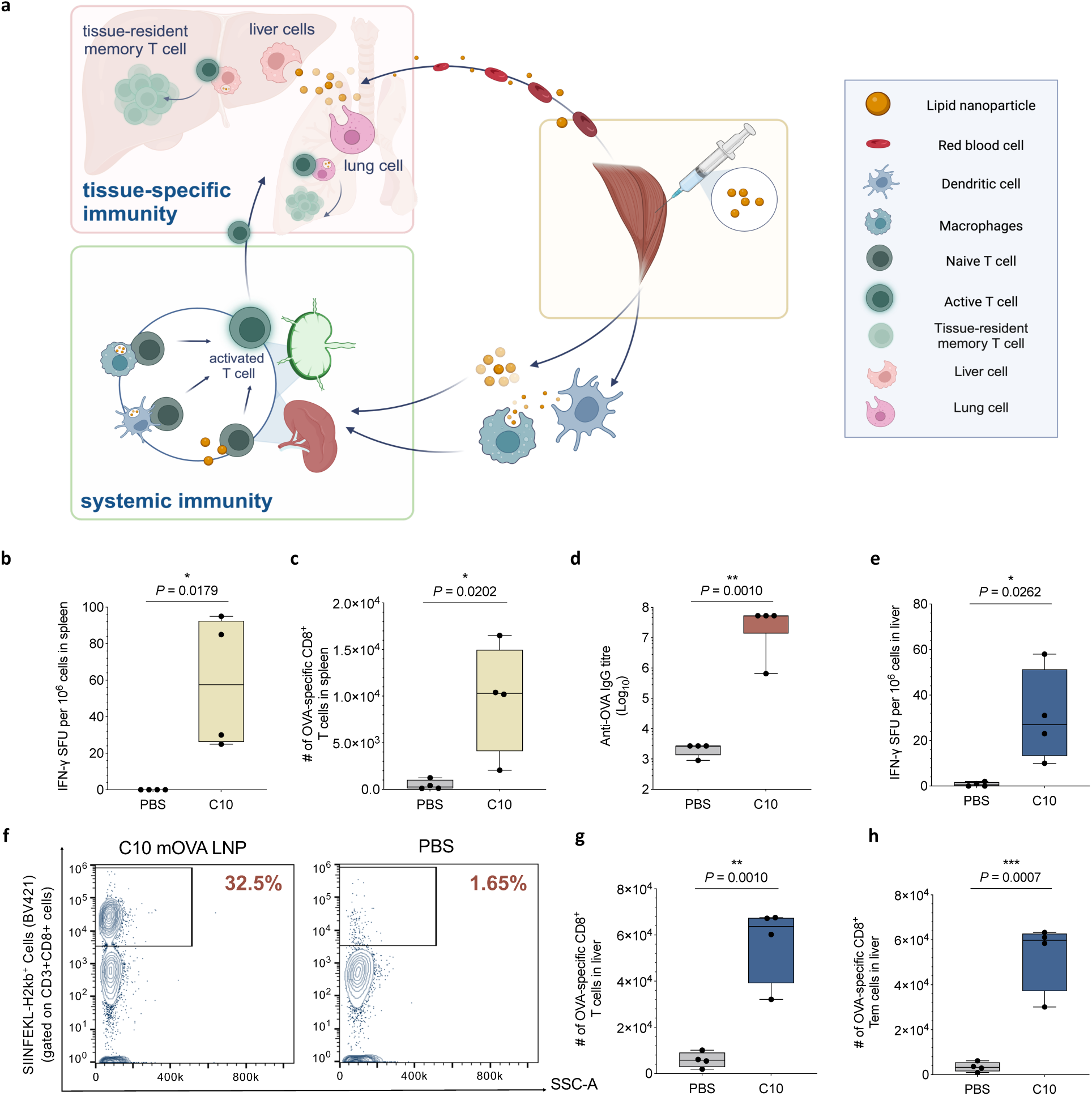
Induction of tissue-specific T cell responses following i.m. injection of mRNA LNPs. **a**, Schematic illustration of LNP trafficking following i.m. injection. **b**, C57BL/6 mice were given three i.m. injections, three weeks apart, of PBS or C10 LNPs loaded with mOVA (10 µg mOVA per injection). Mice were sacrificed four weeks after the final injection, and their lymphocytes were isolated from spleens and livers for analysis. Splenocytes were restimulated *in vitro* with SIINFEKL peptide (2 µg/mL SIINFEKL) and assessed via ELISpot to determine the frequency of IFN-γ-producing cells in spot-forming unit (SFU). **c**, Cellaca MX High-throughput Automated Cell Counter (HACC) was employed to count the lymphocytes within the harvested tissues. Unstimulated splenocytes were assessed via flow cytometry to determine the count of cells positive for CD3, CD8, and SIINFEKL-H-2Kb tetramer. The gating strategy is shown in **Supplementary Fig. 2**. **d,** Titres of OVA-specific serum IgG on day 70, determined by ELISA. **e**, Lymphocytes isolated from the liver were restimulated *in vitro* and assessed via ELISpot to determine the SFU frequency according to the same method described above. **f**–**h**, Representative flow cytometry plots for determining OVA-specific CD8^+^ T cells in the liver **(f)** are shown. Unstimulated lymphocytes isolated from the liver were assessed via flow cytometry to determine the count of cells positive for CD3, CD8, and SIINFEKL-H-2Kb tetramer **(g)**, as well as cells for high CD44 and low CD62L **(h)**. Data are from *n* = 4 biologically independent samples. “C10” stands for C10 mOVA LNPs. Data were analyzed using an unpaired *t*-test. For box plots, the box extends from the 25^th^ to the 75^th^ percentiles with whiskers depicting the minimum/maximum, and the line in the middle of the box is plotted at the median. \**P* < 0.05, \*\**P* < 0.01, \*\*\**P* < 0.001, \*\*\*\**P* < 0.0001.

## Results

### Tissue-specific T cell responses following i.m. injection of mRNA LNPs

Given the limited understanding of mRNA LNP vaccine-mediated T cell responses at the tissue level, we first aimed to examine both systemic and tissue-specific immune responses induced by an LNP formulation (C10 mRNA LNP) recently developed for anti-tumor immunity through systemic Th1 and Th2 immune modulation^34^, with a particular focus on the CD8^+^ T cell responses in the liver. C57BL/6 mice were administered C10 LNPs containing 10 µg ovalbumin mRNA (mOVA) by i.m. injection in the right quadriceps muscle on Days 0, 21, and 42. On Day 70, we observed a significant increase in the number of IFN-γ-secreting lymphocytes (*P* = 0.0179) isolated from the spleens of LNP-treated mice compared to the controls (**Fig. 1b, Supplementary Fig. 1**). Additionally, a marked elevation in the number of OVA-specific CD8^+^ T cells was detected in the spleens for the LNP-treated group, showing a 21-fold increase compared with the PBS control (**Fig. 1c**). Moreover, the C10 mOVA LNPs induced a significantly higher anti-OVA IgG titer (average of 1:10^7^, *P* = 0.0010) than the PBS group, indicating a strong humoral response (**Fig. 1d**). These data confirm that C10 mOVA LNPs can elicit potent and sustained systemic immunity over an extended vaccination period, providing prolonged immune protection.

We next evaluated tissue-specific immunity induced by the C10 LNPs following i.m. injections. Interestingly, in addition to the systemic immunity, a notable increase in IFN-γ-secreting lymphocytes (*P* = 0.0262) and a 10-fold higher frequency of OVA-specific CD8^+^ T cells were detected in the liver of LNP-treated mice compared to the PBS group (**Fig. 1e–g**). Furthermore, a substantial population of CD8^+^ effector memory T (T_EM_) cells (average of 5 × 10^4^, *P* = 0.0007 compared to the PBS group) was observed in the liver, indicating the potential of CD8^+^ T_RM_ cell formation (**Fig. 1h**). Previous studies have predominantly reported local transfection at the injection site and systemic immunity, with limited focus on liver-specific immune responses following i.m. delivery^34^. In this experiment, a tissue-specific T cell response was observed in a non-lymphoid organ, suggesting the potential of using these T cells for rapid and effective immune responses at the critical site of pathogen entry. We then further investigated the mechanism underlying the generation of local immunity following i.m. administration of mRNA LNP vaccines.

### Systemic LNP trafficking results in tissue-specific antigen expression and subsequent T-cell recruitment

To examine the trafficking profile of LNPs following i.m. injection, we administered 10 µg of Cy-5-labelled C10 mRNA LNPs into the right quadriceps of C57BL/6 mice and monitored their biodistribution at 12 and 24 h post-injection. At both time points, LNPs were detected in the injection site and major organs, indicating trafficking from the injection site to the circulation (**Fig. 2a,b**). Notably, at the 24-h time point, most LNPs (51.7%) that travelled out of the injection site were found in the liver, suggesting a biased trafficking pattern.

**Fig. 2:**
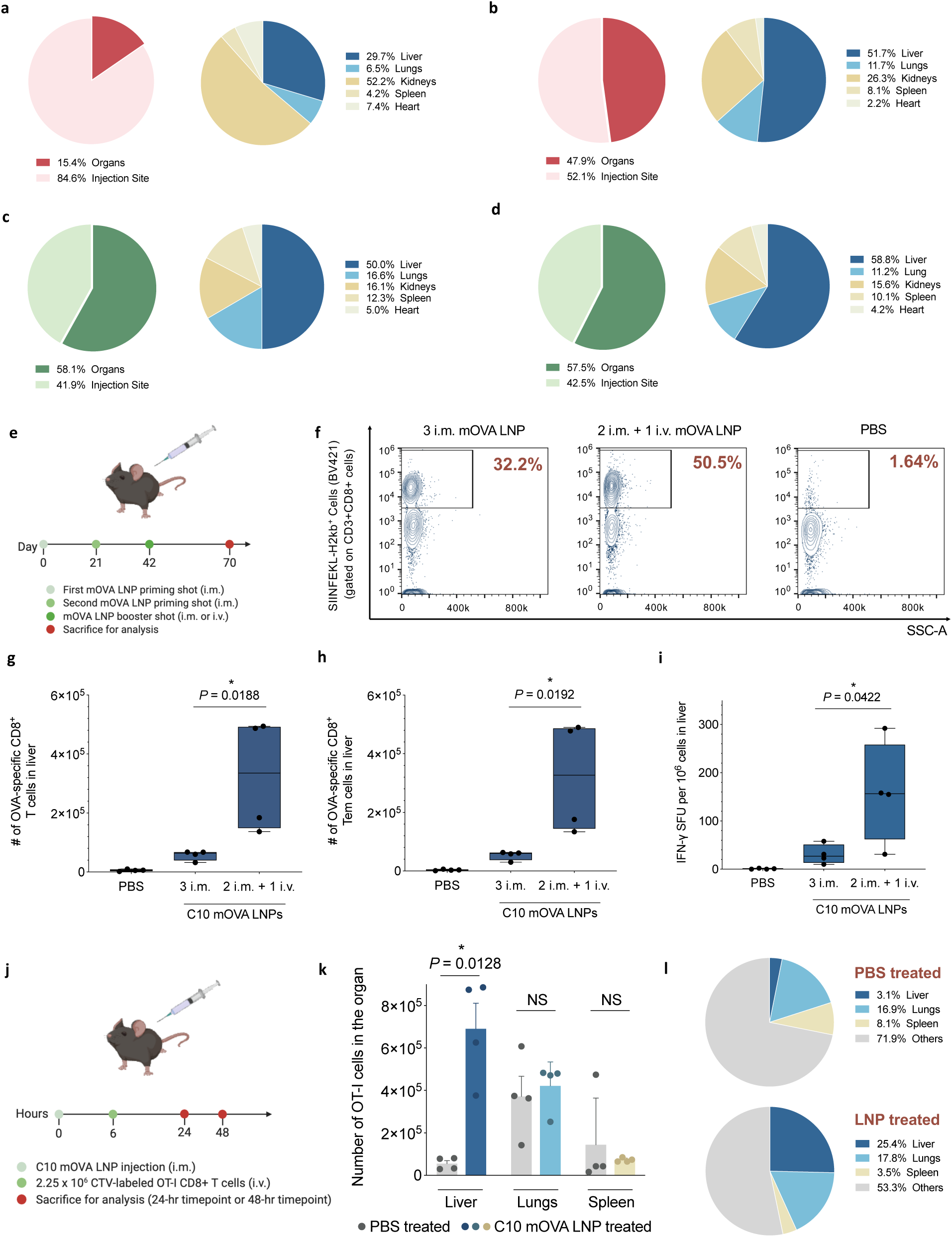
Assessment of mRNA LNP trafficking following i.m. injection and the subsequent immune responses. **a,b**, Biodistribution at 12 h **(a)** and 24 h **(b)** post i.m. injection of C10 LNPs (10 µg Cy5-labelled mRNA per mouse) in C57BL/6 mice, assessed via *ex vivo* fluorescence imaging with IVIS. Data are expressed as the percentage of total radiant efficiency at the injection site versus organs, as well as the percentage distribution across individual organs (n = 4). **c,d**, Total bioluminescence flux in injection site and organs at 24 h **(c)** and 48 h **(d)** post i.m. injection of C10 mLuc LNPs (10 µg mLuc per mouse) in C57BL/6 mice, assessed via *ex vivo* bioluminescence imaging with IVIS. Data are expressed as the percentage of total flux at the injection site versus organs, as well as the percentage expression across individual organs (n = 4). **e**, Timeline for the immune activation experiment. C57BL/6 mice were given three injections (3 i.m. or 2 i.m. + 1 i.v.), three weeks apart, of PBS or C10 LNPs loaded with mOVA (10 µg mOVA per injection). Mice were sacrificed four weeks after the final injection, and their lymphocytes were isolated from spleens and livers for analysis. **f–h**, The Cellaca MX HACC was employed to count the lymphocytes within the harvested tissues. Representative flow cytometry plots for determining OVA-specific CD8^+^ T cells in the liver **(f)** are shown. Unstimulated lymphocytes isolated from the liver were assessed via flow cytometry to determine the count of cells positive for CD3, CD8, and SIINFEKL-H-2Kb tetramer **(g)**, as well as cells for high CD44 and low CD62L **(h)**. The gating strategy is shown in **Supplementary Fig. 2**. **i**, Lymphocytes isolated from the liver were restimulated *in vitro* and assessed for the SFU frequency. **j**, Timeline for OT-I cell migration experiment. CD8^+^ T cells were isolated from OT-I mice and labeled with CellTrace™ Violet (CTV). C57BL/6 mice were first given one i.m. injection of PBS or C10 LNPs loaded with mOVA (10 µg mOVA per mouse). Six hours post-injection, the same mice were given one i.v. injection (tail vein) of 2.25 × 10^6^ CTV-labelled OT-I CD8^+^ T cells. Mice were sacrificed 24 or 48 h post LNP/PBS injection, and their cells were isolated from spleens, livers, and lungs for analysis. **k**, The Cellaca MX HACC was employed to count the lymphocytes within the harvested tissues. Isolated cells were assessed via flow cytometry to determine the count of cells positive for CD3, CD8, and CTV in each organ at 48 h post LNP injection. **l**, Biodistribution of CTV-labeled OT-I cells across three major organs in PBS-treated and LNP-treated mice at 48 h. Percentages represent the proportion of OT-I cells from the total number originally injected (2.25×10^6^). Data are from n = 4 biologically independent samples. Data were analyzed using one-way ANOVA and Tukey’s multiple comparisons test for **g–i**, and an unpaired t-test for **k**. For box plots, the box extends from the 25^th^ to the 75^th^ percentiles with whiskers depicting the minimum/maximum, and the line in the middle of the box is plotted at the median. For bar plots, data represent mean ± s.e.m. NS: *P* > 0.05, \**P* < 0.05, \*\**P* < 0.01, \*\*\**P* < 0.001, \*\*\*\**P* < 0.0001.

To determine if LNP transfection occurred at the sites where they trafficked, C57BL/6 mice were injected with 10 µg of firefly luciferase-encoding mRNA (mLuc) LNPs intramuscularly, and the luciferase protein expression levels at the injection site and in major organs were measured at 24 and 48 h post-injection. As shown in **Fig. 2c,d**, a substantial portion of luciferase expression was found in major organs (58.1%) in addition to the local injected muscle (41.9%). Among the transfected tissues, liver accounted for 50.0% and 58.8% of expression at 24-h and 48-h time points, respectively, consistent with the biodistribution profile. Thus, i.m.-injected C10 LNPs trafficked in part to the liver and result in considerable mRNA-encoded protein expression in the liver.

To validate the causal relationship between LNP trafficking and formation of liver-specific immunity, the third i.m. injection in the vaccination schedule was replaced with an i.v. injection to directly introduce LNPs into the bloodstream and further enhance liver expression (**Fig. 2e**). As illustrated in **Fig. 2f–i**, mice that received an i.v. injection of C10 mOVA LNPs as the third dose exhibited a 6.0-fold increase in the numbers of CD8^+^ T cells and T_EM_ cells, as well as a 5.2-fold increase in the frequency of IFN-γ-secreting lymphocytes in the liver, compared to mice that received three doses by i.m. injection. These data collectively suggest that a higher degree of antigen expression in the liver leads to an elevated level of tissue-specific cellular immune response, supporting the role of LNP trafficking following i.m. injection in generating localized immune responses within the liver.

Our results showed that a substantial portion of LNPs can traffic to the liver after i.m. injection, mediate antigen expression, and enriched antigen-specific CD8^+^ T cells including CD8^+^ T_RM_ cells in the liver. To further elucidate the connection between LNP-mediated transfection and the recruitment of immune cells, we explored if the accumulation of OVA-specific T cells in the liver was driven by tissue-specific antigen expression. We injected and monitored migration of OT-I cells 6 h after i.m. injection of LNPs. OT-I cells are CD8^+^ T cells expressing a transgenic T cell receptor specific for the SIINFEKL peptide epitope. C57BL/6 mice were first given (i.m.) C10 LNPs containing 10 µg of mOVA. Six hours post-injection, 2.25×10^6^ CellTrace Violet (CTV)-labeled OT-I cells were introduced into mice i.v., and the distribution of OT-I cells was subsequently monitored at the 24-h and 48-h time points (**Fig. 2j**). At the 24-h time point, the difference in OT-I cell migration was minor between LNP-treated and PBS-treated groups, with a slight increase in the number of OT-I cells in the liver of the LNP-treated mice (**Supplementary Fig. 3**). However, by 48 h, there was a more than 6.0-fold elevation (*P* = 0.01) in the number of OT-I cells in the liver of LNP-treated mice compared to the PBS-treated group, while the number of OT-I cells in other organs remained unchanged (**Fig. 2k, Supplementary Fig. 4**). As depicted in **Fig. 2l**, only 3.1% of the total number of injected OT-I cells were detected in the liver in the PBS-treated group. In contrast, the liver had the highest percentage of OT-I cells among the three organs examined in the LNP-treated group, accounting for 25.4% of the total injected OT-I cells. These findings suggest that T cells are being recruited to the liver, presumably in response to liver-specific expression of antigen induced by mRNA LNPs that trafficked from the i.m. injection site to the liver.

### LNP trafficking and transfection of major organs is LNP composition-dependent

We next determined whether LNP trafficking and transfection in major organs vary among different LNP formulations. We selected three LNP compositions based on FDA-approved formulations: MDN (used in the Moderna COVID-19 vaccine^41^), PBT (using the ALC-0315 as the ionizable lipid and same composition as in the Pfizer-BioNTech COVID-19 vaccine^42^, except DMG-PEG2000 as the PEGylated lipid), and ALN (used in the Alnylam Onpattro formulation^43^), in addition to one LNP formulation DI-8, which was described in our previous report showing high levels of gene expression in the liver through i.v. injections^22^.

The biodistribution profiles of these selected LNP formulations were first evaluated. C57BL/6 mice received one i.m. injection of LNPs containing 10 µg of Cy-5-labelled mRNA at their right quadriceps; and the biodistribution of the LNPs was monitored at 12-h and 24-h time points (**Fig 3a**). Trafficking was observed beyond the injection site for all formulations. As depicted in **Fig. 3a**, distinct trafficking patterns were observed amongst the selected formulations. For instance, the DI-8 exhibited 13.9% “leakage” from the injection site into the systemic circulation, whereas the MDN LNPs yielded a much higher leakage of 31.4% (*P* < 0.0001). Despite the differences in leakage, once the LNPs entered the bloodstream, they displayed comparable biodistribution patterns across major organs with ∼40% to the liver, ∼10% to the lungs, and ∼30% to the spleen (**Fig. 3b, Supplementary Figs. 5 and 6**).

**Fig. 3:**
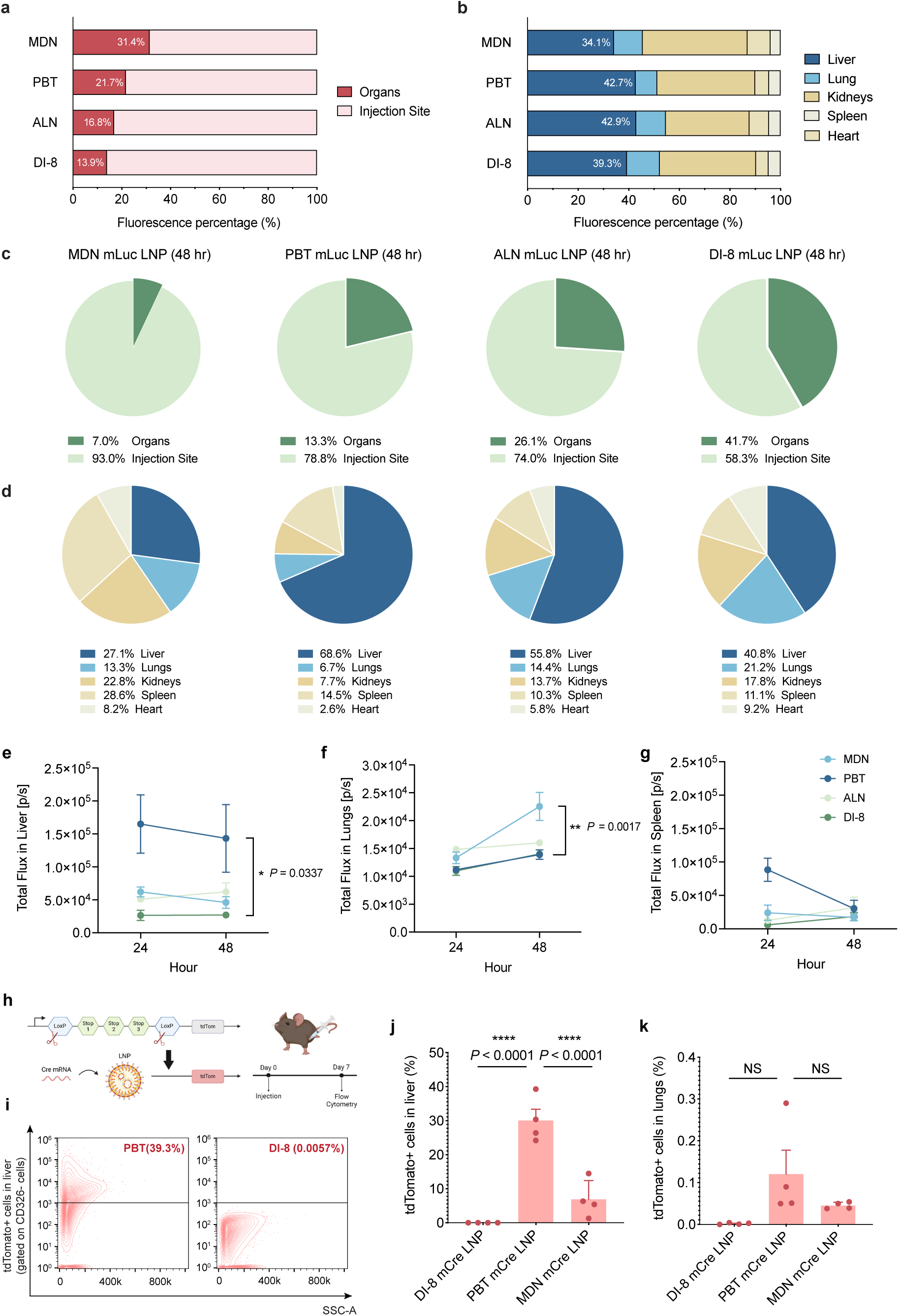
Systemic trafficking of different LNP formulations following i.m. injection. **a,b**, Biodistribution at 24 h post i.m. injection of four selected LNPs (10 µg Cy5-labelled mRNA per mouse) in C57BL/6 mice, assessed via ex vivo fluorescence imaging with IVIS. Data are expressed as the percentage of total radiant efficiency at the injection site versus major organs **(a)**, as well as the percentage distribution across individual organs **(b)**. **c,d**, Total bioluminescence flux at the injection site and organs 48 h post i.m. injection of four selected LNPs containing 10 µg mLuc per dose in C57BL/6 mice, assessed via *ex vivo* bioluminescence imaging using IVIS. Data are expressed as the percentage of total flux at the injection site versus major organs **(c)**, as well as the percentage expression across individual organs **(d)**. **e–g**, Total flux in photons per second generated by each of the four selected LNPs in the liver **(e)**, lungs **(f)**, and spleen **(g)** 24 and 48 h post i.m. injection are shown. **h**, Schematic illustration of Ai9 mouse model used for organ-specific transfection validation and timeline of the Ai9 transfection validation experiment. C57BL/6 mice were given one i.m. injection of selected LNPs loaded 30 µg mCre per mouse. Mice were sacrificed on day 7 post-injection for analysis. **i–k**, The tdTom^+^ cells were quantified via flow cytometry analysis of the cells isolated from the liver and lungs. Representative flow cytometry plots for tdTom^+^ hepatocytes in the liver of PBT and DI-8 groups at 7 days post-injection are shown **(i)**. Percentages of tdTom^+^ cells in the liver **(j)** and lungs **(k)** are shown. Gating strategies are shown in **Supplementary Figs. 8 and 9**. Data are from n = 4 biologically independent samples. Data were analyzed using one-way ANOVA and Tukey’s multiple comparisons test for **j,k**. For bar plots, data represent mean ± s.e.m. NS: *P* > 0.05, \**P* < 0.05, \*\**P* < 0.01, \*\*\**P* < 0.001, \*\*\*\**P* < 0.0001.

Next, we evaluated the transfection efficiency of the selected LNP formulations in different tissues. Each C57BL/6 mouse received one i.m. injection of LNPs containing 10 µg of mLuc at the right quadriceps; and the luciferase expression levels at the injection site and the major organs were measured at 24- and 48-h post-injection. As depicted in **Fig. 3c**, different LNP formulations exhibited distinct transfection patterns outside of the injection site. Despite similar biodistribution patterns following systemic trafficking, these LNPs showed different transfections in major organs, independent of the biodistribution pattern (**Fig. 3d**). The total luminescence intensity measurements revealed that the PBT LNPs achieved the highest level of mLuc expression in the liver (*P* = 0.0377 compared with DI-8). Meanwhile, the MDN LNPs showed the highest level of mLuc expression in the lungs compared to the other formulations (*P* = 0.0017 compared with DI-8). On the other hand, the DI-8 showed minimal transfection levels in the liver and lungs, with the most transfection localized at the injection site (**Fig. 3e–g**). Although these LNP compositions share similar physical properties such as size, PDI, zeta potential, and encapsulation efficiency (**Supplementary Fig. 7, Supplementary Table 2**), they exhibited different tissue-specific transfection profiles following systemic trafficking from the i.m. injection site. Based on these results, we selected PBT, MDN, and DI-8 LNPs as representative formulations with biased tissue-specific expressions in the liver, lungs, and no systemic organ transfection, respectively.

To further assess the organ-specific trafficking and transfection, we gave one i.m. injection of Cre-recombinase mRNA (mCre) LNPs to genetically engineered tdTomato (tdTom) reporter (Ai9) mice that contain a LoxP-flanked stop cassette that prevents expression of the tdTom protein. This mouse model enables the identification of transfected cells through Cre-recombinase expression, which removes the stop cassette, allowing the fluorescent tdTom protein to be expressed (**Fig. 3h**). Seven days following i.m. injection, the PBT LNP formulation resulted in the highest percentage, 39.3% (*P* < 0.0001) of tdTomato-positive (tdTom^+^) cells in the liver compared to the MDN LNPs (5.4%) and DI-8 LNPs (0.01%) (**Fig. 3i,j, Supplementary Fig. 10**). In the lungs, there were no significant differences in transfection efficiency across the three LNP formulations, as all three exhibited relatively low levels of tdTom^+^ cells (**Fig. 3k, Supplementary Fig. 10**).

To explore whether injection volume will alter the extent and distribution of the LNP trafficking following i.m. injection, we evaluated the gene expression efficiency in major organs with different i.m. injection volumes. As shown in **Supplementary Figs. 11 and 12**, changing the i.m. injection volume from 50 to 100 µL did not affect the percentage of tdTom^+^ hepatocytes in the liver, indicating that the observed LNP trafficking was driven by intrinsic LNP properties rather than diffusion kinetics that may be influenced by the injection volume **(Supplementary Figs. 11 and 12**). This reaffirms the importance of LNP composition in modulating the extent and pattern of the LNP trafficking outside the injection site for organ-targeted applications.

### Assessment of short-term systemic and tissue-specific immune activation by selected mRNA LNP formulations following i.m. injection

We next investigated the effect of biased LNP trafficking and expression in different organs on the generation of tissue-specific immunity. We evaluated short-term immune responses following a prime-and-boost vaccination regimen using the PBT, MDN, and DI-8 LNP formulations, with OVA as the model antigen (**Fig. 4a**). On Day 28, the frequency of OVA-specific CD8^+^ T cells in the liver of mice treated with PBT mOVA LNPs was 4.3-, 8.4-, and 28.5-fold higher (*P* < 0.0001) than that of mice treated with the MDN mOVA LNPs, DI-8 mOVA LNPs, and PBS, respectively (**Fig. 4b,c**). A substantial population of OVA-specific CD8^+^CD44^hi^CD62L^lo^CD69^hi^ T_RM_-like cells was detected in the liver for the PBT mOVA LNP-treated group, with the number being 4.6-, 3.9-, and 22.7-fold higher (*P* = 0.0002, *P* = 0.0001, and *P* < 0.0001, respectively) than the MDN mOVA LNP-, DI-8 mOVA LNP-, and PBS-treated groups, respectively (**Fig. 4d**). This indicates successful induction of tissue-specific immunity and establishment of T_RM_ cells in the liver with the PBT LNPs, which are crucial for long-term protection. In the lungs, there were 2.3-, 9.6-, and 48.3-fold increases (*P* = 0.0064, *P* < 0.0001, and *P* < 0.0001, respectively) in OVA-specific CD8^+^ T cell frequencies after treatment with MDN mOVA LNPs, compared with PBT mOVA LNPs, DI-8 mOVA LNPs, and PBS, respectively (**Fig. 4e,f**). The number of OVA-specific CD8^+^CD44^hi^CD62L^lo^CD69^hi^ T_RM_-like cells in the lungs was also 2.7-, 7.4, and 9.4-fold higher (*P* = 0.0316, *P* = 0.0021, and *P* = 0.0015, respectively) in the MDN mOVA LNP-immunized group in comparison to those immunized with PBT mOVA LNPs, DI-8 mOVA LNPs, and PBS, respectively (**Fig. 4g**). This demonstrates that MDN mOVA LNPs elicited stronger lung-specific CD8+ T cell immunity compared to the other LNP formulations.

**Fig. 4:**
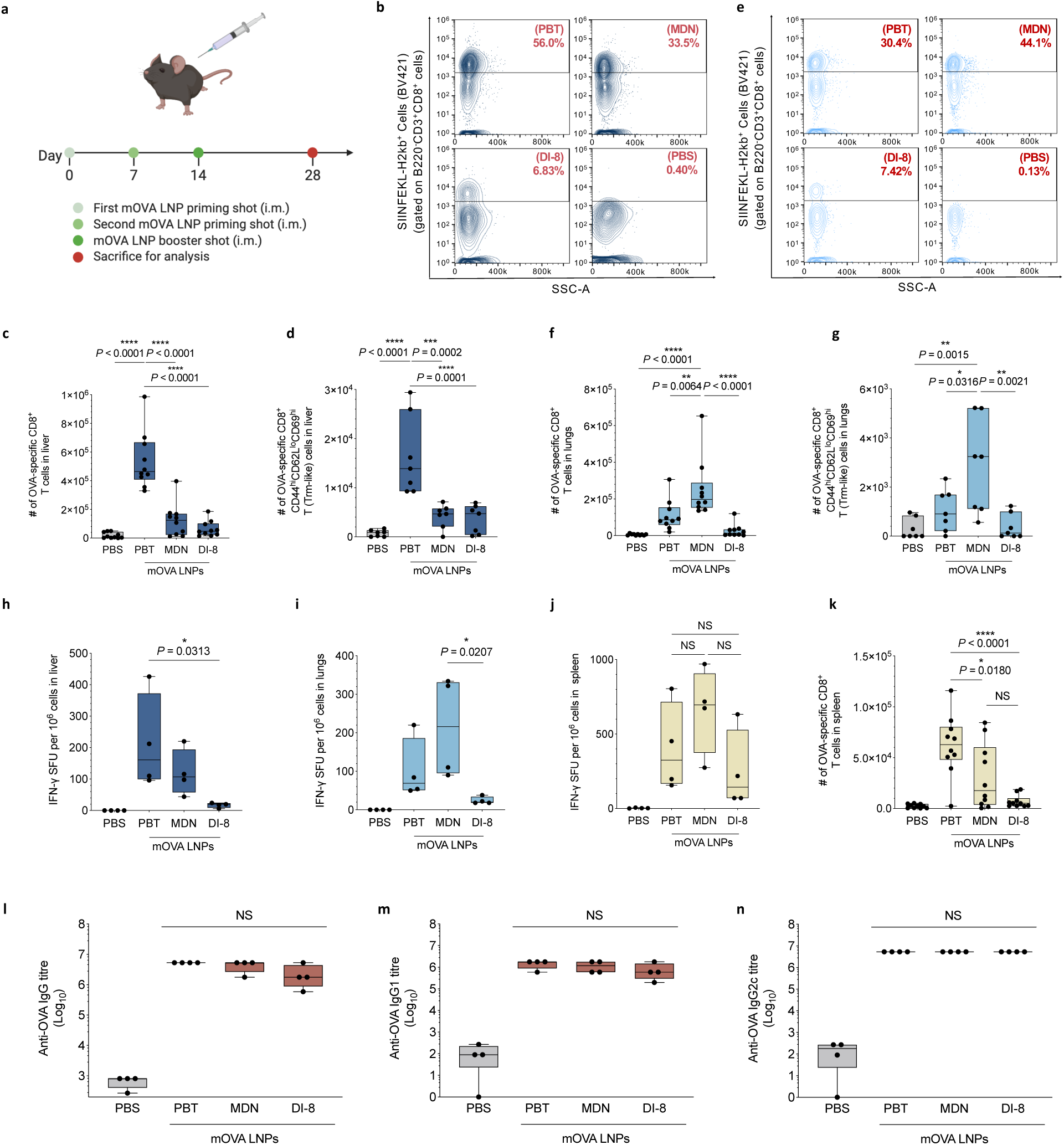
Short-term immune assessments of selected mRNA LNPs following i.m. injection. **a**, Timeline for the short-term vaccination study. C57BL/6 mice were given three i.m. injections, one week apart, of PBT, MDN, or DI-8 LNPs containing 10 µg mOVA per injection or PBS. Mice were sacrificed two weeks after the final injection, and their lymphocytes were isolated from the liver, lungs, and spleen for analysis. **b–d**, The Cellaca MX HACC was employed to count the isolated lymphocytes within the harvested tissues. Representative flow cytometry plots for determining OVA-specific CD8^+^ T cells in the liver **(b)** are shown. Unstimulated lymphocytes isolated from the liver were assessed via flow cytometry to determine the count of cells positive for CD3, CD8, and SIINFEKL-H-2Kb tetramer **(c)**, as well as high CD44, low CD62L, and high CD69 **(d)**. The gating strategy is shown in **Supplementary Fig. 13**. **e–g**, Representative flow cytometry plots for determining OVA-specific CD8^+^ T cells in the lungs **(e)** are shown. Unstimulated lymphocytes isolated from the lungs were assessed via flow cytometry to determine the count of cells positive for CD3, CD8, and SIINFEKL-H-2Kb tetramer **(f)**, as well as high CD44, low CD62L, and high CD69 **(g)**. **h–j**, Lymphocytes isolated from the liver, lungs, and spleen were restimulated *in vitro* with SIINFEKL peptide (2 µg/mL SIINFEKL) and assessed via FluoroSpot to determine the SFU frequency in the liver **(h)**, lungs **(i)**, and spleen **(j)**. **k**, Number of cells positive for CD3, CD8, and SIINFEKL-H-2Kb tetramer in the spleen, assessed via flow cytometry. **l–n**, Titres of OVA-specific IgG, IgG1, and IgG2c antibodies in blood serum on day 28, determined by ELISA. Data represent n = 10 biologically independent samples from three independent experiments (**b,e,c,f,k**), n = 7 biologically independent samples from two independent experiments (**d,g**), and n = 4 biologically independent samples from one representative experiment (**h–j,l– n**). Data were analyzed using one-way ANOVA and Tukey’s multiple comparisons test for **c–d** and **f–n**. For box plots, the box extends from the 25^th^ to the 75^th^ percentiles with whiskers depicting the minimum/maximum, and the line in the middle of the box is plotted at the median. NS: *P* > 0.05, \**P* < 0.05, \*\**P* < 0.01, \*\*\**P* < 0.001, \*\*\*\**P* < 0.0001.

To further assess the antigen-specific cytolytic functionality of the lymphocytes, the liver, lungs, and spleen of the vaccinated mice were collected on Day 28 and homogenized into a cell suspension for *ex vivo* antigen restimulation. Elevated production of pro-inflammatory cytokine after restimulation was detected across all three LNP formulations. Particularly, the PBT mOVA LNP-treated group showed a significantly greater number of IFN-γ-secreting lymphocytes in the liver, with the frequency being 1.8- and 12.4-fold higher compared to the MDN mOVA LNP- and DI-8 mOVA LNP-treated groups, confirming the superior liver-biased immune response generation capability of the PBT mOVA LNPs (**Fig. 4h, Supplementary Fig. 14**). In contrast, the MDN formulation induced the highest number of IFN-γ-secreting lymphocytes in the lungs, reflecting its lung-bias characteristic (**Fig. 4i, Supplementary Fig. 15**). In the spleen, the number of IFN-γ-secreting lymphocytes was similar across all three LNP formulations, demonstrating comparable systemic cellular immune responses (**Fig. 4j, Supplementary Fig. 16**). However, the frequency of OVA-specific CD8^+^ T cells, specifically, in the spleen for the PBT mOVA LNP formulation is 2.0- and 8.8-fold higher than that for the MDN and DI-8 LNP formulations, respectively, suggesting potential for more OVA-specific CD8^+^ T cells in systemic circulation and subsequent homing to antigen-expressing tissues (**Fig. 4k**). In terms of humoral immunity, all three formulations elicited similar levels of OVA-specific IgG, IgG1, and IgG2c titers indicative of potent systemic humoral responses (**Fig. 4l–n**). These results are consistent with the expectation that systemic humoral immunity is primarily driven by local transgene transfection and CD4^+^ helper T cell pathways, and less on the organ-specific trafficking of the LNPs.

Collectively, these data provide evidence that different LNP compositions can lead to marked differences in the magnitude and development of tissue-specific immunity within specific organs.

### Assessment of long-term systemic and tissue-specific immune activation by selected mRNA LNP formulations following i.m. injection

Next, we assessed the longevity of the immune responses and memory formation generated by the selected LNP formulations to determine whether the local and systemic immunity observed at the earlier time point would be sustained over an extended period. To evaluate long-term immune responses, we followed the same prime-and-boost vaccination schedule with three i.m. injections of LNPs containing 10 µg of mOVA or PBS. On Day 90, the liver, lungs, spleen, and blood of the mice were collected for immune assessments (**Fig. 5a**). In line with the 28-day short-term immune response data, the PBT LNP formulation continued to show a substantial retention of OVA-specific CD8^+^ T cells (12.7- and 64.9-fold more than the MDN and DI-8 LNP formulations, respectively) and OVA-specific CD8^+^CD44^hi^CD62L^lo^CD69^hi^ T_RM_-like cells (18.7- and 64.0-fold more than the MDN and DI-8 LNP formulations, respectively) in the liver. Notably, the number of these cells remained substantially higher than PBS baseline and the other treatment groups, with the difference becoming even more pronounced at this extended time point than the 28-day time point. As illustrated, the MDN and DI-8 LNP formulations showed minimal retention of OVA-specific CD8^+^ T cells and T_RM_-like cells in the liver, showing levels close to the PBS baseline (**Fig. 5b–d**). In the lungs, a modest but discernible increase in the number of OVA-specific CD8^+^ T cells (1.5- and 2.1-fold more than the PBT and DI-8 LNP formulations, respectively) and OVA-specific CD8^+^CD44^hi^CD62L^lo^CD69^hi^ T_RM_-like cells (1.6- and 3.4-fold more than the PBT and DI-8 LNP formulations, respectively) were observed in the MDN group at the 90-day mark, suggesting that MDN may favor the formation of longer-term immune memory in lung tissues (**Fig. 5e–g**). However, these frequencies remained lower than those observed in the liver with the PBT formulation, highlighting the differences in targeting capability and room for further optimization for lung-targeting LNPs.

**Fig. 5:**
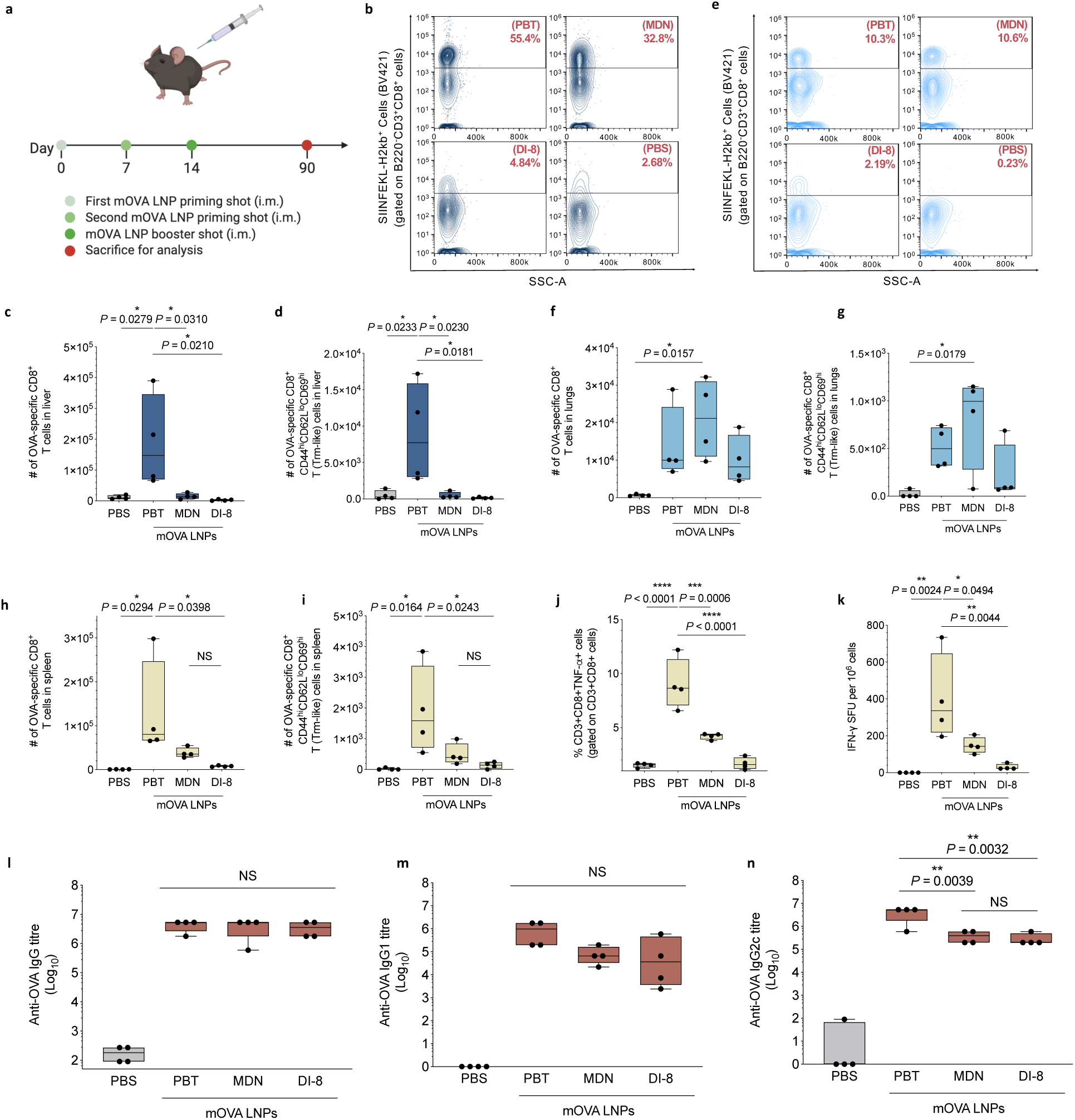
Long-term immune assessments of selected mRNA LNPs following i.m. injection. **a**, Timeline for the long-term vaccination study. C57BL/6 mice were given three i.m. injections, one week apart, of PBT, MDN, or DI-8 LNPs containing 10 µg mOVA per injection or PBS. Mice were sacrificed 2.5 months after the final injection, and lymphocytes were isolated from the liver, lungs, and spleen for analysis. **b–d**, The Cellaca MX HACC was employed to count the isolated lymphocytes within the harvested tissues. Representative flow cytometry plots for determining OVA-specific CD8^+^ T cells in the liver **(b)** are shown. Unstimulated lymphocytes isolated from the liver were assessed via flow cytometry to determine the count of cells positive for CD3, CD8, and SIINFEKL-H-2Kb tetramer **(c)**, as well as high CD44, low CD62L, and high CD69 **(d)**. The gating strategy is shown in **Supplementary Fig. 13**. **e–g**, Representative flow cytometry plots for determining OVA-specific CD8^+^ T cells in the lungs **(e)** are shown. Unstimulated lymphocytes isolated from the lungs were assessed via flow cytometry to determine the count of cells positive for CD3, CD8, and SIINFEKL-H-2Kb tetramer **(f)**, as well as high CD44, low CD62L, and high CD69 **(g)**. **h–i**, Unstimulated lymphocytes isolated from the spleen were assessed via flow cytometry to determine the count of cells positive for CD3, CD8, and SIINFEKL-H-2Kb tetramer **(h)**, as well as high CD44, low CD62L, and high CD69 **(i)**. **j**, Lymphocytes isolated from the spleen were restimulated *in vitro* with OVA and SIINFEKL peptide (100 μg/mL OVA and 2 μg/mL SIINFEKL) for 12 h and assessed via intracellular cytokine staining and flow cytometry and to determine the percentages of CD3^+^CD8^+^TNFα^+^ cells. **k**, Lymphocytes isolated from the spleen were restimulated *in vitro* with SIINFEKL peptide (2 µg/mL SIINFEKL) and assessed via FluoroSpot to determine the SFU frequency. **l–n**, Titres of OVA-specific IgG, IgG1, and IgG2c antibodies in blood serum on day 90 post-vaccination, determined by ELISA. Data are from representative experiments with n = 4 biologically independent samples. Data were analyzed using one-way ANOVA and Tukey’s multiple comparisons test for **c–d** and **f–n**. For box plots, the box extends from the 25^th^ to the 75^th^ percentiles with whiskers depicting the minimum/maximum, and the line in the middle of the box is plotted at the median. NS: *P* > 0.05, \**P* < 0.05, \*\**P* < 0.01, \*\*\**P* < 0.001, \*\*\*\**P* < 0.0001.

Interestingly, at the 90-day time point, in the spleen of mice vaccinated with the PBT LNPs, a significantly higher number of OVA-specific CD8^+^ T cells (3.4- and 17.2-fold more than the MDN and DI-8 LNPs, respectively) and CD8^+^CD44^hi^CD62L^lo^CD69^hi^ T_RM_-like cells (3.9- and 14.2-fold more than the MDN and DI-8 LNPs, respectively) was observed (**Fig. 5h,i**). To assess the antigen-specific cytolytic T cell responses, the spleen of the vaccinated mice were collected and homogenized into a cell suspension for *ex vivo* antigen restimulation. The three LNP-treated groups showed increased frequencies of CD3^+^CD8^+^TNF-α^+^ cell population compared to the PBS group. In comparison to MDN and DI-8 LNPs, the PBT LNPs resulted in a significantly higher frequency of CD3^+^CD8^+^TNF-α^+^ cells with *P* values of 0.0006 and <0.0001, respectively (**Fig. 5j, Supplementary Fig. 17**). Additionally, the frequency of IFN-γ-secreting lymphocytes in the spleen of PBT LNP-treated mice was 2.7- and 13.1-fold higher compared to MDN and DI-8 LNP-treated mice (**Fig. 5k, Supplementary Fig. 18**). We hypothesize that the biodistribution and transfection of mRNA LNPs in organs outside of the injection site may correlate with the ability of enhancing and maintaining systemic immunity.

Regarding humoral immunity, the anti-OVA IgG and IgG1 titers detected at Day 90 were comparable across all three formulations, indicating that general antibody responses were relatively stable (**Fig. 5l,m**). However, the PBT LNPs induced a significantly higher level of IgG2c titer than the MDN and DI-8 LNPs with *P* values of 0.0039 and 0.0032, respectively, pointing to a more potent Th1-biased immune response (**Fig. 5n**). This finding is consistent with our earlier data, which showed a sustained systemic effector T cell response, further reinforcing the role of the PBT LNP composition in promoting a durable Th1-driven immune response.

Taken together, these results provide evidence that the PBT LNP formulation not only supports robust immunity in the liver, but also enhances long-term systemic immune memory, particularly through retention of CD8^+^ T cells in the spleen and promotion of a Th1-biased systemic immune profile. This finding indicates that LNPs differ substantially in their ability to induce both tissue-specific and systemic immune responses critical for long-lasting vaccine-mediated protection.

### Proof-of-concept therapeutic and prophylactic efficacy of mRNA LNP vaccine with liver-biased T cell recruitment after i.m. administration

Building on the localized antigen-specific CD8^+^ T cell responses in the liver elicited by the PBT LNPs, we next evaluated its therapeutic potential as a cancer vaccine against liver tumor growth using OVA as the model antigen in mice. To address the challenge of accurately measuring tumor size within the liver, B16-OVA-fLuc cells, which are B16 melanoma cells genetically engineered to constitutively express luciferase, were utilized to enable non-invasive monitoring of tumor growth via bioluminescence imaging. First, C57BL/6 mice were inoculated with 5×10^6^ B16-OVA-fLuc cells via intrahepatic injection on Day 0. On Days 2, 4, and 6 post-tumor inoculation, the mice were immunized with PBT, MDN, or DI-8 LNPs containing 10 µg mOVA (**Fig. 6a**). Over the 30-day observation period, mice treated with PBT LNPs showed prolonged overall survival (**Fig. 6b**) and considerable tumor regression (**Fig. 6c**) in the B16-OVA-fLuc treatment model; their 30-day survival rate was 57% compared with median survival times of 6 days, 4 days, and 5 days for MDN LNP, DI-8 LNP, and PBS groups, respectively. In the PBT group, there was apparent tumor regression from Days 5 to 30 (**Fig. 6d**).

**Fig. 6:**
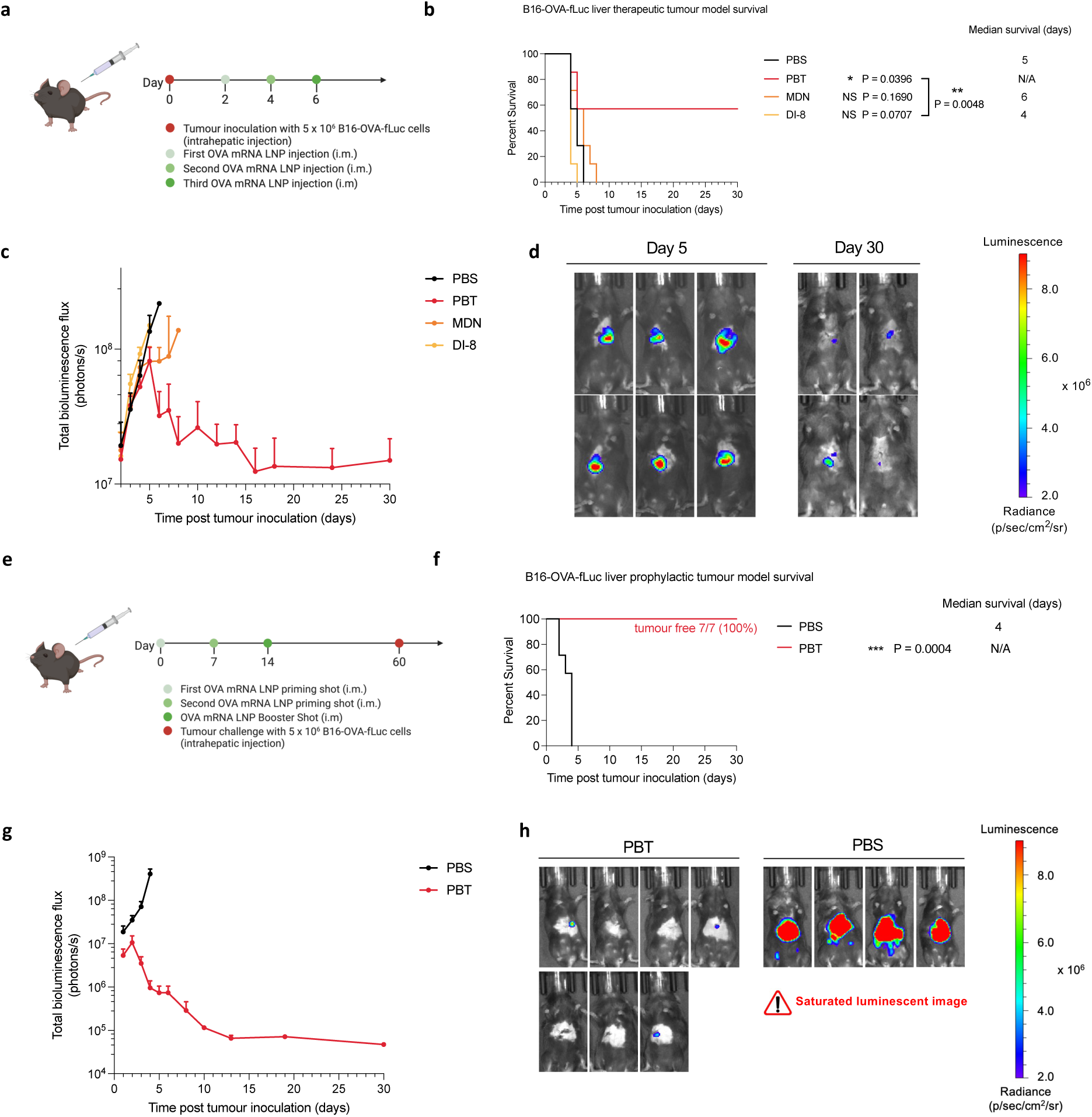
Antitumor efficacy of the mRNA LNP formulations as therapeutic and prophylactic vaccines. **a–d**, Schematic and results of a therapeutic vaccination model for B16-fLuc-OVA in C57BL/6 mice. Mice were inoculated via intrahepatic injection with B16-fLuc-OVA cells and then given three i.m. injections, 2 days apart, of PBT, MDN, or DI-8 LNPs containing 10 µg mOVA per mouse per injection or PBS **(a)**. Survival curves **(b)**, tumor luminescence **(c)**, and representative IVIS images of the PBT group on days 5 and 30 **(d)** are shown. **e–g**, Schematic and results of a prophylactic vaccination model for B16-fLuc-OVA in C57BL/6 mice. Mice were given three i.m. injections of LNPs containing 10 µg mOVA per injection or PBS, 1 week apart before intrahepatic inoculation of B16-OVA-fLuc cells on Day 60 **(e)**. Survival curves **(f)**, tumor luminescence **(g)**, and all IVIS images on Day 4 **(h)** are shown. “PBT,” “MDN,” and “DI-8” stand for PBT mOVA LNPs, MDN mOVA LNPs, and DI-8 mOVA LNPs, respectively. Data represent mean ± s.e.m. with n = 7 biologically independent samples. Survival curves were compared using the log-rank Mantel-Cox test. NS: *P* > 0.05, \**P* < 0.05, \*\**P* < 0.01, \*\*\**P* < 0.001, \*\*\*\**P* < 0.0001.

We next evaluated the efficacy of PBT LNPs as a prophylactic vaccine in the B16-OVA-fLuc mouse liver tumor model. C57BL/6 mice were immunized on Days 0, 7, and 14 with 10 µg of PBT LNPs containing 10 µg mOVA. On Day 60, these mice were challenged with 5 × 10^6^ B16-OVA-fLuc cells by intrahepatic injection (**Fig. 6e**). As illustrated in **Fig. 6f,g**, PBT-vaccinated mice had 100% survival rate during the 30-day observation period, compared to a median survival of 4 days for the PBS group. Although the PBT-vaccinated mice initially displayed a detectable increase in tumor burden, it was quickly controlled and reduced to near-background levels for all vaccinated mice. This highlights the ability of the PBT LNP formulation to elicit a protective immune response capable of controlling and eradicating tumor cells even after a delayed challenge.

By directing LNPs to traffic and express the antigen in specific organs affected by metastatic tumors, it is possible to deliver immunotherapeutic agents directly to the tumor microenvironment. This tissue-targeted delivery and generation of T_RM_-like cells can enhance the local immune response against tumor cells, improving the efficacy of cancer immunotherapy.

## Discussion

The systemic LNP trafficking behavior following i.m. and s.c. injections has been noted in previous literature^37–40,44^. However, these discussions have been predominantly confined to the role of lymphatic drainage, with limited exploration into the concurrent entry of LNPs into the systemic circulation for therapeutic implications. Despite reports showing the trafficking of mRNA LNPs into other organs, attention has been on potential adverse effects due to the “leakage” rather than investigating their therapeutic potential^37–40^. Furthermore, the relationship between LNP formulation composition and trafficking profiles, including distribution to and transgene expression in the liver and lungs following i.m. injection, has remained largely unexplored, particularly regarding the associated potential therapeutic benefits. This study explores the utility of systemic trafficking of the LNPs for therapeutic benefits. By comparing the biodistribution and transfection profiles of different LNP formulations, we identified those with distinct organ-biased expression properties, particularly the PBT LNPs showing a substantial liver-biased trafficking and antigen expression following i.m. injection. The PBT LNPs trafficked to and transfected the liver tissue, resulting in localized antigen expression, which, in turn, promoted the recruitment of antigen-specific T cells and enhanced T_RM_-like cell retention within the liver over 90 days. As a result, this LNP formulation demonstrated sustained local immunity and prolonged T cell effector function. Additionally, we observed that the humoral immune response was less dependent on systemic LNP trafficking. Despite differences in the extent of LNP trafficking from the injection site, the three selected LNP formulations examined in this study elicited comparable humoral immunity against the antigen of interest. This suggests that systemic LNP trafficking primarily influences T cell-mediated immunity, particularly in shaping antigen-specific CD8^+^ T cell responses and CD8^+^ T_RM_ cell formation, while having minimal influence on mobilizing humoral immune responses. The dissociation between systemic LNP trafficking and humoral immunity highlights the distinct mechanisms governing B cell and T cell activation and recruitment in response to LNP-delivered antigens. These findings provide additional insights into how tissue-specific T-cell responses induced by LNP trafficking from the injection site can promote long-term immunity, with important implications for both localized and systemic defense mechanisms.

Previous research on LNPs has underscored the effect of LNP composition on tissue-targeting and transfection capabilities with i.v. injections^26–32,22,33,34^. Our findings expand on this by demonstrating that organ-specific LNP trafficking following i.m. injection can be similarly influenced by LNP composition. In this study, changing LNP formulations led to varying degrees of systemic trafficking. The ability to tailor LNP composition for enhanced and targeted systemic trafficking may open new possibilities for creating LNPs with controllable trafficking profiles for therapeutic purposes. One notable observation was that formulations with higher leakage capabilities also have higher PEGylate lipid content (**Supplementary Table. 2**). The hydrophilic nature of PEG contributes to reduced aggregation and longer circulation times^45^ and possibly reduced binding with the extracellular matrix, which may allow for more efficient escape from the injection site and subsequent tissue distribution, offering a potential explanation for higher systemic trafficking out of the injection site. Further studies are needed to elucidate the exact mechanism by which LNP composition controls the systemic trafficking and tissue-specific antigen expression following i.m. injection, which could guide the development of more effective and targeted LNP-based therapeutic vaccines.

Tissue-specific CD8^+^ T cells, particularly CD8^+^ T_RM_ cells, have gained attention for their role in long-lasting protection in tissues as entry points for pathogens^1–4^. These cells are recruited following antigen expression in local tissues, with some establishing long-term residence to deliver localized, rapid immune responses upon antigen re-exposure^5–12^. In various infectious disease studies, T_RM_ cells have been shown to play a critical role in protection against infection, highlighting their potential in therapeutic strategies^1,4,8,10,46,47^. By inducing tissue-specific T cell immunity beyond systemic immune responses, robust and durable protection can be conferred in tissues where pathogens may enter and/or reside, offering a promising dual-protective strategy in vaccine development. Our study illustrates the utility of this approach in preventing tumor recurrence, leveraging both systemic and local immune responses to enhance overall efficacy.

In this proof-of-concept mouse liver tumor study, we identified that the PBT LNP formulation, with more efficient trafficking to and transgene expression in the liver following i.m. injection, exhibited the strongest capacity for controlling and suppressing liver tumor engraftment, extending survival in both therapeutic and prophylactic models. This LNP formulation effectively mobilized immune defenses to prevent tumor recurrence by engaging tissue-specific CD8^+^ T cells including T_RM_ cells while sustaining systemic immunity. This approach is particularly valuable for preventing tumor recurrence after surgical excision by maintaining immune surveillance in the tissue where the tumor initially resided. Additionally, it may help prevent tumor metastasis to critical organs, reducing the risk of secondary tumor formation. From a tumor immunotherapy perspective, modulating LNP trafficking to target tumor-residing tissues offers a more effective strategy for tumor control and clearance. Furthermore, the principles demonstrated in this tumor model could be applied to diseases like malaria, where targeting the liver during the latency period in hepatocytes could prevent disease progression. This highlights the broader potential of LNP-based vaccines to address both infectious diseases and cancers by tailoring immune responses to specific tissues.

In summary, our study provides insights into systemic trafficking of LNPs from the injection site following i.m. injection and revealed that mRNA LNP compositions exhibiting a higher propensity for systemic entry following i.m. injection and subsequent antigen expression in selected organs, induced localized antigen-specific CD8^+^ T cell response and the formation of T_RM_-like cells. Using a tumor vaccine as a proof-of-concept case study, we showed that the PBT mRNA LNPs were capable of generating antigen expression and T cell recruitment to the liver beyond the systemic immunity following i.m. immunization, resulting in enhanced antitumor efficacy in both therapeutic and prophylactic liver tumor models. These findings have important implications for the design of targeted vaccines and therapeutics where precise tissue-specific and durable immunity are key. The ability to finetune LNP composition to achieve desired leakage profiles through i.m. injection not only enhances therapeutic efficacy, but also opens new possibilities for creating highly specific and effective treatments for a range of diseases.

## Methods

### Materials

DLin-MC3-DMA was obtained from MedKoo Biosciences, and SM-102 and ALC-0315 were from Broadpharm. DSPC, DOPE, 18PG, and DMG-PEG-2000 were from Avanti Polar Lipids, and cholesterol was from Sigma-Aldrich. The B16-OVA-fLuc cells, a luciferase reporter cell line, were obtained from AcceGen. All mRNA constructs were purchased from TriLink BioTechnologies, capped using their CleanCap proprietary co-transcriptional capping method, and designed form the naturally occurring Cap 1 structure with a high efficiency. The mRNAs were also polyadenylated, modified with 5-methoxyuridines, and optimized for mammalian systems. D-Luciferin (sodium salt) was acquired from Gold Biotechnology.

### LNP synthesis and characterization

LNPs were made by mixing an organic phase consisting of the lipid components and an aqueous phase consisting of the gene cargo. The organic phase was prepared by dissolving a mixture of ionizable lipid (ALC-0315, SM-102, or DLin-MC3 DMA), cholesterol, DMG-PEG2000, and a helper lipid (DOPE, DSPC, or 18PG) at predetermined ratios in ethanol. The mRNA (OVA mRNA, fLuc mRNA, or Cre mRNA) was dissolved in 25 mM magnesium acetate buffer at a pH of 4.0. The flash nanocomplexation (FNC) device was used to mix the ethanol and aqueous phases at a 3:1 ratio using syringe pumps using a previously reported protocol^48^. Then the LNPs were purified through dialysis against DI water with a 100-kDa MWCO cassette at 4 °C for 24 h and stored at 4 °C before injection. The zeta potential, average size, and polydispersity index (PDI) of LNPs were measured using dynamic light scattering (ZetaPALS, Brookhaven Instruments) method, with diameters reported as the intensity average.

A Quant-iT RiboGreen assay (ThermoFisher, R11490) was utilized to quantify mRNA concentrations and assess the encapsulation efficiency (EE) of LNP formulations. Standard curves were prepared in duplicate on a black 96-well plate with mRNA concentrations ranging from 0.1 to 1.0 µg/mL in each well. Samples of LNPs were added in triplicate at a consistent LNP to DI water ratio of 1:20. Following this, 60 µL of RiboGreen solution—a 200-fold dilution of RiboGreen dye in water—was added to each well. Fluorescence measurements were taken using a BioTek Synergy H1 plate reader (excitation: 480 nm, emission: 520 nm) to determine mRNA levels in intact LNPs. Following the initial reading, 10% w/v Triton-X was added to each well to disrupt the LNP structure and release the encapsulated mRNA. Similar to the first reading, 60 µL of RiboGreen solution was added to the wells. Fluorescence readings were measured to determine the total mRNA content post-release. The measured fluorescence signals before and after mRNA release were converted to mRNA concentration using the standard curves. The encapsulation efficiency (EE%) of the LNPs were calculated using the following equation.

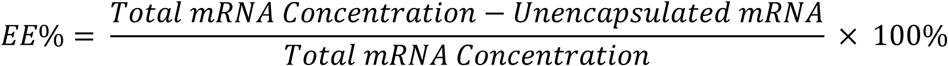

### Animals and primary cells

All animal procedures were conducted in accordance with protocols approved by the Johns Hopkins Institutional Animal Care and Use Committee (Protocol #MO24E165). C57BL/6 mice (male and female), aged 6–8 weeks, were obtained from the Jackson Laboratory. Ai9 mice (male and female), 6–8 weeks old, were bred in the Johns Hopkins Animal Facilities and assigned randomly to groups in the studies. Similarly, male and female OT-I mice, aged 6–8 weeks, were bred in the same facilities and randomly grouped. The mice had free access to pelleted feed and water, with the feed typically containing 5% fiber, 20% protein, and 5–10% fat. On average, the mice consumed 4–5 g of pelleted feed (120 g per kg body weight) and drank 3–5 mL of water (150 mL per kg body weight) daily. The temperature in the mouse rooms was maintained between 18– 26 °C (64–79 °F) with 30–70% relative humidity, ensuring at least 10 air changes per hour. The mice were housed in standard shoebox cages with corncob bedding.

The LNPs were given through i.m. (right quadriceps) injection or i.v. (lateral tail vein) injection at a predetermined dose per mouse. The LNP suspensions were concentrated to 200 μg/mL for i.m. injection or 100 μg/mL for i.v. injection of mRNA by an Amicon Ultra-2 centrifugal filter unit with an MWCO of 100 kDa. The D-luciferin solution was given through i.p. injection (lower quadrant of abdomen) at a predetermined dose per mouse. For experiments in Ai9 mice, the Cre mRNA LNP formulations were prepared as described above and administered via i.m. or i.v. injections at an mRNA dose of 10 μg per mouse. For experiments described in **Supplementary Fig. 12–13**, the LNP suspensions were concentrated to 100 μg/mL and 200 μg/mL for the 100 μL and 50 μL injection groups.

For the OT-I cell migration experiments, primary CD8 T cells were obtained from homogenized spleens and inguinal lymph nodes of OT-I mice using a CD8^+^ T cell isolation kit and LS columns (Miltenyi Biotec), following the manufacturer’s protocols. Briefly, splenocytes were obtained from the homogenized suspension by lysing red blood cells with the ACK lysis buffer (ThermoFisher Scientific) and incubated with the supplied antibody cocktail and magnetic beads, then passed through a magnetic column to obtain a purified population. Isolated CD8^+^ T cells were stained with CellTrace Violet cell proliferation kit (ThermoFisher, C34571). Cell counts were adjusted to 2.25 × 10^6^ per 100 μL in PBS, and mice were injected i.v. (lateral tail vein) with 100 μL of cell suspension.

### Tissue processing and cell isolation

For isolation of cells from the liver, lungs, and spleen in the Ai9 mouse experiments, the harvested tissues were placed onto 40-μm cell strainers and digested mechanically with the back of a 3-mL syringe plunger in PBS. The cells were pelleted at 300 ×g for 5 min at 4 °C, followed by resuspension in the ACK lysis buffer and incubation at room temperature for 7 min to lyse red blood cells. Cells were then pelleted by centrifugation at 300 ×g for 5 min at 4 °C, washed with PBS and pelleted twice before staining for flow cytometry. All steps were performed protected from light.

For isolation of lymphocytes from the liver, harvested livers were placed onto 40-μm cell strainers. The livers were gently dissociated by pressing through the strainer with the back of a 3-mL syringe plunger in circular or back-and-forth motions, and the resulting cell suspensions were collected in sterile 50-mL tubes. The cell strainer and plunger were rinsed with 15 mL of RPMI-1640 media, followed by an additional 15-mL rinse, for a total of 30 mL RPMI-1640 media. The tubes containing the liver cell suspension were swirled to ensure even distribution and kept on ice. To pellet gross hepatocytes, tubes were centrifuged at 77 ×g for 1 min at 4 °C without brake. The supernatant containing lymphocytes was transferred to fresh 50-mL tubes, and the volume was adjusted to 50 mL with RPMI-1640 media. Cells were pelleted by centrifugation at 478 ×g for 8 min at 4 °C with brake. After aspirating the supernatants, the cell pellets in each tube were resuspended in 10 mL of the 35% Percoll solution (Millipore-Sigma), prepared in HBSS/Heparin solution. Cell suspensions were then centrifuged at 850 ×g for 25 min at room temperature without brake. Following centrifugation, the top layer containing hepatocytes and most of the Percoll solution was carefully aspirated. The lymphocyte pellets were resuspended in 2 mL of ACK lysis buffer and incubated at room temperature for 5 min to lyse red blood cells. Eight mL of MACS buffer was then added, and the cells were pelleted by centrifugation at 417 ×g for 8 min at 4 °C with brake. Isolated lymphocytes were suspended in RPMI-1640 media for subsequent analyses.

For isolation of lymphocytes from the spleen, harvested spleens were placed onto 40-μm cell strainers. The spleens were mechanically digested through the cell strainers with the back of a 3-mL syringe plunger in a lymphocyte separation medium (PromoCell). The resulting single-cell cell suspensions were transferred to sterile 15-mL tubes, and 1 mL of RPMI-1640 media was slowly added to each tube to form visible layers. The tubes were then centrifuged at 800 ×g for 25 min at 4 °C without brake. Following centrifugation, the middle layer containing lymphocytes was carefully collected and transferred to fresh 15-mL tubes, and 5 mL of RPMI-1640 media was added and mixed thoroughly. The cells were pelleted by centrifugation at 300 ×g for 5 min at 4 °C. The pellet was then resuspended in 3 mL of RPMI-1640 media and centrifuged again at 300 ×g for 5 min at 4 °C. Isolated lymphocytes were suspended in RPMI-1640 media for subsequent analyses.

For isolation of lymphocytes from the lung, harvested lungs were first minced into smaller pieces using surgical scissors and transferred to sterile 50-mL tubes. Lung digestion solution was prepared with RPMI-1640 and collagenase type 1 (45 U/µL collagenase I). Ten mL of lung digestion solution was added to each tube containing the minced lung tissue, and the samples were incubated on a shaker at 37 °C for 1 h. The digested samples were then further processed mechanically with the back of a 3-mL syringe plunger through 40-μm cell strainers. RPMI-1640 medium was used to wash cells through the filters into sterile 50-mL tubes. The cell suspensions were then centrifuged at 500 ×g for 5 min at 4 °C. After centrifugation, the cell pellets were resuspended in 2 mL of ACK lysis buffer and incubated at room temperature for 5 min to lyse red blood cells. Eight mL of MACS buffer was then added, and the cells were pelleted by centrifugation at 500 ×g for 5 min at 4 °C with brake. Isolated lymphocytes were suspended in RPMI-1640 media for subsequent analyses.

### Antibodies and staining for flow cytometry

Antibody panels are provided in **Supplementary Table 1**. All antibodies were diluted at a ratio of 1:100 before use. LIVE/DEAD fixable dead cell stain kits were used to determine the viability of cells. CellTrace Violet cell proliferation kit (ThermoFisher, C34571) was used to stain and monitor OT-I cells *in vivo*. eBioscience Foxp3/Transcription Factor Staining buffer set (ThermoFisher, 00-5523-00) was used for intracellular staining.

Isolated cells from the tissues, as described in the previous section, were resuspended and pelleted in 100 µL of antibodies diluted in flow cytometry staining buffer obtained from eBioscience™. The cells were then incubated on ice in the dark for 1 h. After the incubation period, the stained cells were washed twice with PBS and subsequently resuspended in 200 µL of eBioscience™ flow cytometry staining buffer for flow cytometry analysis. Flow data was acquired using an Attune NXT flow cytometer and analyzed with FlowJo software v.10.

### Enzyme-linked immunosorbent spot (ELISpot) assay and FluoroSpot assay

For the ELISpot assay, multiscreen filter plates (Millipore-Sigma, S2EM004M99) were coated with antibodies targeting IFN-γ (BD Biosciences, 551881) and subsequently blocked according to the manufacturer’s instructions. A total of 1×10^6^ isolated lymphocytes were then added to each well and stimulated with SIINFEKL peptide at a concentration of 2 μg/mL for 18 h. The spots were detected using a mouse IFN-γ detection antibody (BD Biosciences, 551881), followed by an incubation with streptavidin-HRP (BD Biosciences, 557630) and AEC substrate (BD Biosciences, 551951). The plates were then forwarded to the SKCCC Immune Monitoring Core for further analysis. For the FluoroSpot assays, the Mouse IFN γ FluoroSpot Plus kit (Mabtech) was utilized, adhering to the manufacturer’s protocols. In this case, 5×10^5^ isolated lymphocytes were plated per well and stimulated with SIINFEKL peptide at 2 μg/mL for 18 h before sending the plates to the SKCCC Immune Monitoring Core for evaluation.

### Enzyme-linked immunosorbent assay (ELISA)

For serum antibody detection, 100 μL of blood sample was drawn from the tail vein of immunized mice on the day of sacrifice. Levels of antigen-specific IgG, IgG1, and IgG2c in the serum were measured by ELISA. Flat-bottomed 96-well plates (Nunc) were precoated with OVA protein at a concentration of 2 μg protein per well in 100 mM carbonate buffer (pH 9.6) at 4°C overnight, which were then blocked at room temperature for 2 h with blocking buffer, which consisted of 10% BSA in PBS. The serum obtained from mice was first diluted 30 times in the blocking buffer, followed by threefold serial dilutions. After blocking was done, the plates were washed with PBS-T (PBS containing 0.05% Tween), and the diluted serum samples were then added to the wells and incubated at 4 °C overnight. Horseradish peroxidase-conjugated goat anti-mouse IgG, IgG1, and IgG2c (Southern Biotech Associates) were used at a dilution of 1:2,000, 1:4,000, and 1:4,000, respectively, in the blocking buffer for labeling. After 1-h antibody incubation at room temperature, the plates were washed with PBS-T, followed by incubation with TMB ELISA substrate solution at room temperature. After a 30-min incubation, 50 µL of 4 N sulfuric acid was added to the wells to quench the reaction. The plates were then read at a wavelength of 450 nm with a plate reader. A sample was considered positive if its absorbance was twice as much as or higher than the absorbance of the negative control.

### Immunization and tumor experiments

For immunization, C57BL/6 mice aged 6-8 weeks were i.m. injected in their right quadriceps with different LNPs containing 10 μg of OVA mRNA, as described in the main text. For all vaccination studies, a total of three doses were given. In prophylactic and therapeutic liver tumor studies, mice were injected via intrahepatic injection with 5 × 10^6^ B16-OVA-fLuc cells. In therapeutic studies, vaccinations began on Day 2 after tumor inoculation. Tumor growth was measured three times a week through whole-body imaging using IVIS. The D-luciferin solution was given through i.p. injection (lower quadrant of abdomen) at a predetermined dose per mouse. Mice were euthanized when the total luminescence flux from the tumor exceeded 8 × 10^7^ p/sec/cm^2^/sr.

### Statistics

Unpaired t-tests were performed when comparing two groups. One-way analysis of variance (ANOVA) and Tukey’s multiple comparisons were performed when comparing more than two groups. Survival curves were compared using log-rank Mantel–Cox test. Statistical analysis was performed using Microsoft Excel and Prism 10 (GraphPad). A difference was considered significant if *P* < 0.05 (\**P* < 0.05, \*\**P* < 0.01, \*\*\**P* < 0.001, \*\*\*\**P* < 0.0001).

## Reporting Summary

Further information on research design is available in the Nature Portfolio Reporting Summary attached along with this article.

## Data Availability

The main data supporting the results in this study are available within the paper and its Supplementary Information. The raw and analyzed datasets generated during the study are available for research purposes from the corresponding authors on reasonable request. Source data for the figures are provided with this paper.

## Acknowledgments

We thank Christopher Thoburn from the SKCCC Immune Monitoring Core at Johns Hopkins Medicine for technical support. This work was partially supported by the National Institute of Allergy and Infectious Diseases through a grant U01AI155313 (S.C.M. and H.-Q.M.) and the National Institute of Biomedical Imaging and Bioengineering through a grant P41EB028239 (H.Q.M. and J.P.S.).

## Author contributions

C.W., Y.Z., and H.-Q.M. conceived and designed the study. H.-Q.M., S.C.M., and J.P.S. secured the funding for the study. C.W., Y.Z., X.L., K.D.G., D.Y., X.L., J.C., A.T.C., J.M., Y.S., J.L. and S.L. performed the experiments. C.W., Y.Z., X.L., K.D.G., D.Y., X.L., J.C., A.T.C., J.M., Y.S., S.L., J.P.S., S.C.M., and H.-Q.M. participated in data analysis and interpretation. The manuscript was written by C.W., Y.Z., and H.-Q.M., with revisions made by A.T.C., Y.S., J.P.S., and S.C.M., and input from all the other authors.

## Code availability statement

There is no code used in the paper.

