## Supplementary Information for "Systemic trafficking of mRNA lipid nanoparticle vaccine following intramuscular injection generates potent tissue-specific T cell response"

### Table of Contents

**Supplementary Figure 1 |** Representative images of IFN-γ-secreting cells from the enzyme-linked immunosorbent assay.

**Supplementary Figure 2 |** Gating strategy for flow cytometry plots for T cell responses after vaccination.

**Supplementary Figure 3 |** Assessment of OT-I cell migration at 24 h post LNP injection**.**

**Supplementary Figure 4 |** Representative flow cytometry plots for OT-I migration post LNP injection at the 48-h time point.

**Supplementary Figure 5 |** Biodistribution profiles of the selected LNP formulations following i.m. injection in injection site vs. major organs.

**Supplementary Figure 6 |** Biodistribution profiles of the selected LNP formulations following i.m. injection across different major organs.

**Supplementary Figure 7 |** Z-average size and PDI of the four selected LNP formulations measured by DLS (n = 3) after dialysis.

**Supplementary Figure 8 |** Gating strategy for flow cytometry plots for assessment of liver cell transfection by the selected LNP formulations**.**

**Supplementary Figure 9 |** Gating strategy for flow cytometry plots for assessment of lung cell transfection by the selected LNP formulations.

**Supplementary Figure 10 |** Representative flow cytometry plots for assessment of cell transfection in the liver and lungs by the selected LNP formulations.

**Supplementary Figure 11 |** Percentage of tdTom+ cells in the liver and lungs following LNP injection by different administration routes and volumes.

**Supplementary Figure 12 |** Representative flow cytometry plots for tdTom+ cells in the liver and lungs following LNP injection by different administration routes and volumes.

**Supplementary Figure 13 |** Gating strategy for flow cytometry plots for T cell responses after vaccination.

**Supplementary Figure 14 |** Representative images of IFN-γ-secreting cells in the liver at the short-term time point (Day 28) from the FluoroSpot assay.

**Supplementary Figure 15 |** Representative images of IFN-γ-secreting cells in the lungs at the short-term time point (Day 28) from the FluoroSpot assay.

**Supplementary Figure 16 |** Representative images of IFN-γ-secreting cells in the spleen at the short-term time point (Day 28) from the FluoroSpot assay.

**Supplementary Figure 17 |** Representative flow cytometry plots for analysis of CD3+CD8+TNFα+ cells in the spleen at the long-term time point on Day 90.

**Supplementary Figure 18 |** Representative images of IFN-γ-secreting cells in the spleen at the long-term time point on Day 90 from the FluoroSpot assay.

**Supplementary Table 1 |** Anti-mouse antibodies used in flow cytometry panels. Marker, fluorophore, catalogue number, source, and concentration are indicated.

**Supplementary Table 2 |** Composition details and characterization of the five evaluated LNP formulations.

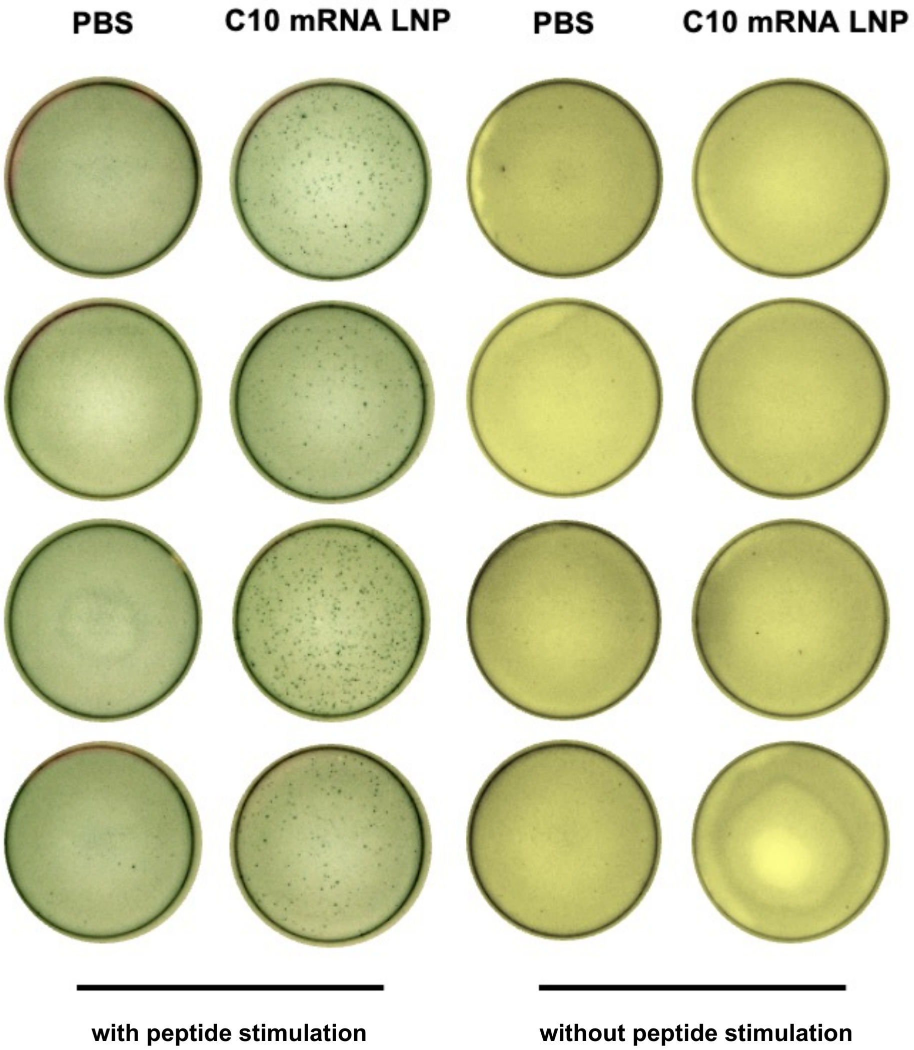


**Supplementary Figure 1. Representative images of IFN-γ-secreting cells from the enzyme-linked immunosorbent assay.** Frequency of IFN-γ-secreting cells among restimulated splenocytes, assessed via ELISpot. Splenocytes were restimulated *in vitro* with SIINFEKL peptide (2 μg/mL SIINFEKL).

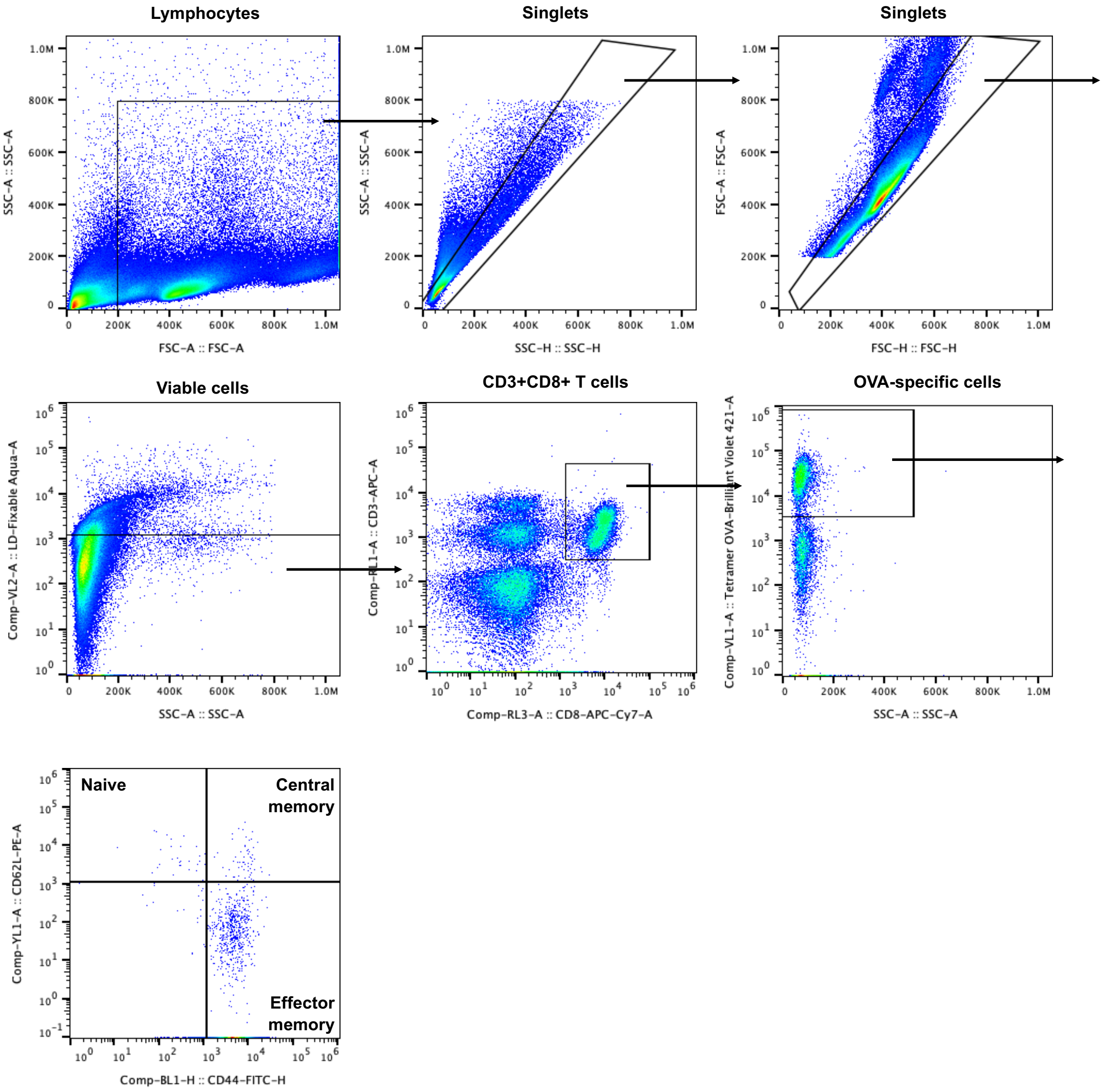


**Supplementary Figure 2. Gating strategy for flow cytometry plots for T cell responses after vaccination.** Initially, lymphocytes isolated from the spleens and livers were selected using SSC-A and FSC-A parameters, followed by singlet selections with SSC-A/SSC-H and FSC-A/FSC-H plots. Viable cells were identified and selected based on the live/dead Fixable Aqua-A and SSC-A plot. Next, the CD3$^+$CD8$^+$ T cell population was selected with downstream analysis focused on antigen (OVA)-specific and CD8$^+$ T cell subtypes.

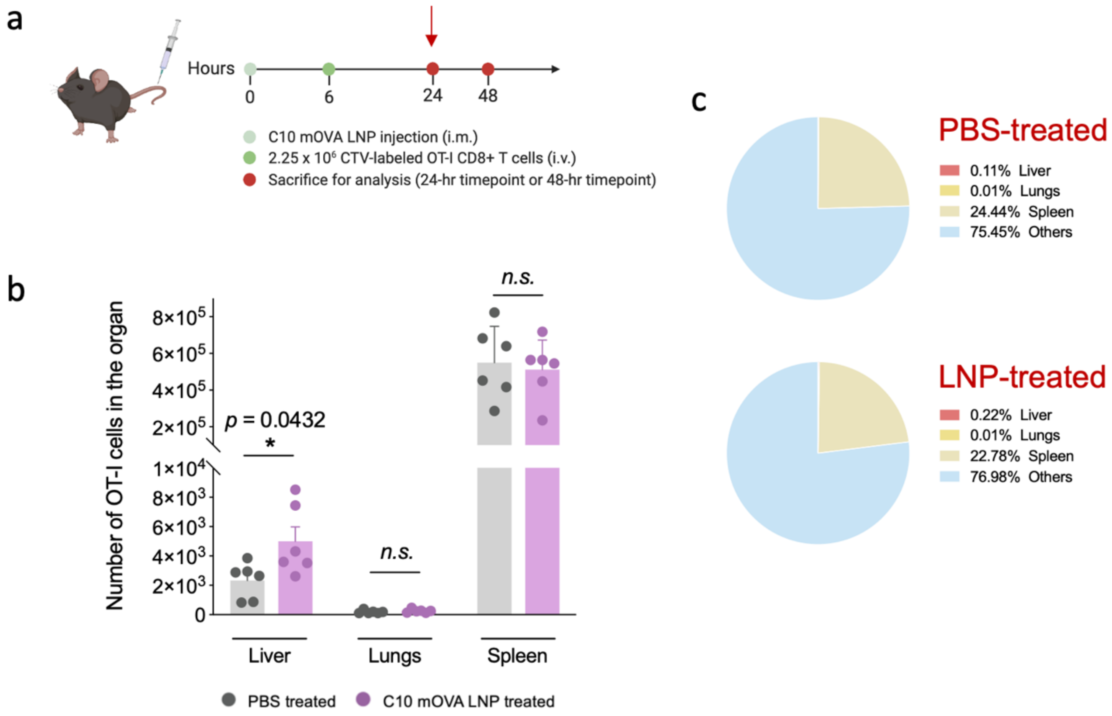


**Supplementary Figure 3. Assessment of OT-I cell migration at 24 h post LNP injection. a,** Timeline for OT-I cell migration experiment. CD8+ T cells were isolated from OT-I mice and labeled with CellTrace™ Violet (CTV). C57BL/6 mice were first given one i.m. injection of PBS or C10 LNPs loaded with mOVA (10 ug mOVA per mouse). Six hours post-injection, the same mice were given one i.v. injection (tail vein) of $2.25 \times 10^6$ CTV-labelled OT-I CD8+ T cells. Mice were sacrificed 24 or 48 hours post LNP/PBS injection, and their cells were isolated from spleens, livers, and lungs for analysis. **b**, The Cellaca MX High-throughput Automated Cell Counter was employed to count the cells within the harvested tissues. Isolated cells were assessed via flow cytometry to determine the count of cells positive for CD3, CD8, and CTV in each organ at 24 h post LNP injection. **c**, Biodistribution of CTV-labeled OT-I cells across three major organs in PBS-treated and LNP-treated mice at the 24-h time point. Percentages represent the proportion of OT-I cells from the total number originally injected ($2.25 \times 10^6$ cells).

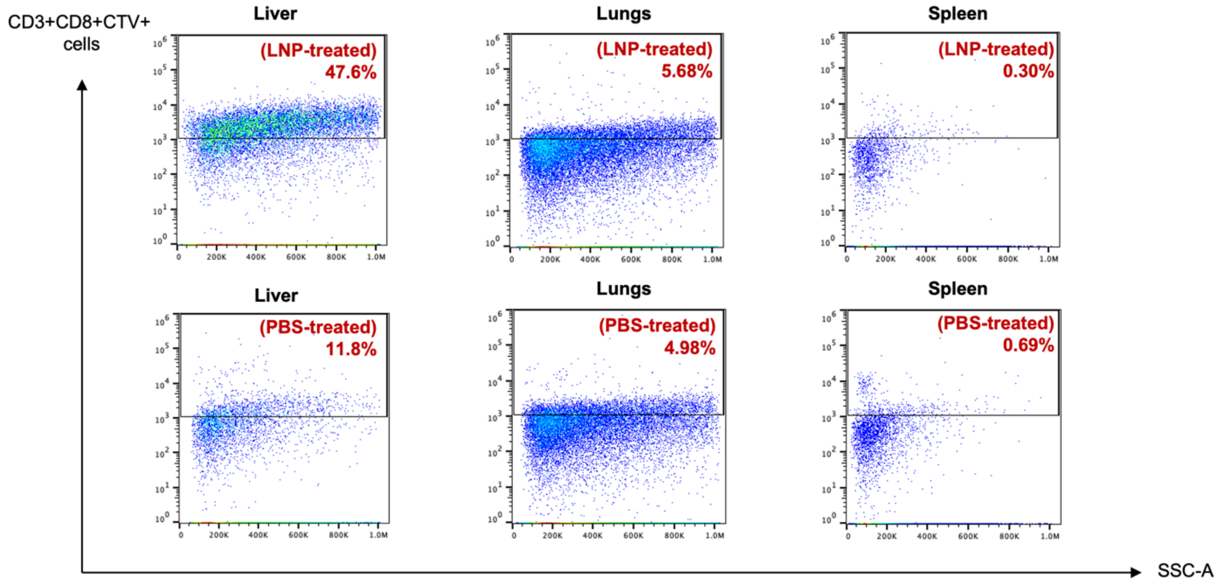


**Supplementary Figure 4. Representative flow cytometry plots for OT-I migration post LNP injection at the 48-h time point.** C57BL/6 mice were first given one i.m. injection of PBS or C10 LNPs loaded with mOVA (10 ug mOVA per mouse). Six hours post-injection, the same mice were given one i.v. injection (tail vein) of $2.25 \times 10^6$ CTV-labelled OT-I $CD8^+$ T cells. Mice were sacrificed 48 h post LNP/PBS injection, and their cells were isolated from the spleen, liver, and lungs for analysis. Percentages of cells positive for CTV gated on $CD8^+$ cells are shown.

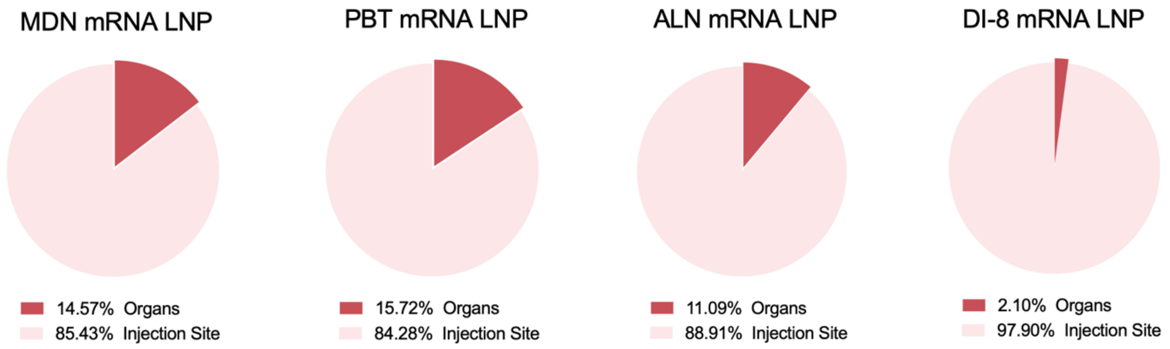


**Supplementary Figure 5. Biodistribution profiles of the selected LNP compositions following i.m. injection in injection site vs. major organs.** Biodistribution at 12 h post i.m. injection of five selected LNPs (10 µg Cy5-labelled mRNA per mouse) in C57BL/6 mice, assessed via *ex vivo* fluorescence imaging with IVIS. Data are expressed as the percentage of total radiant efficiency at the injection site versus major organs.

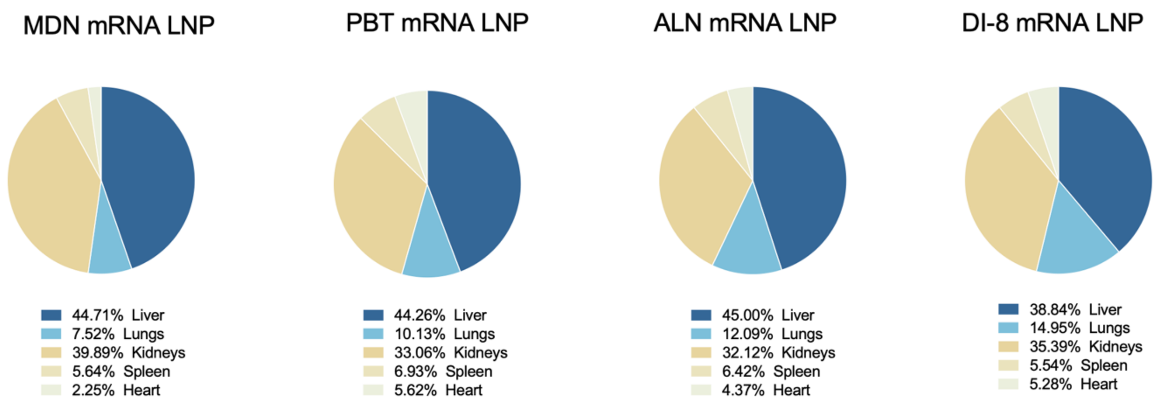


**Supplementary Figure 6. Biodistribution profiles of the selected LNP compositions following i.m. injection across individual organs.** Biodistribution at 12 h post i.m. injection of five selected LNPs (10 µg Cy5-labelled mRNA per mouse) in C57BL/6 mice, assessed via *ex vivo* fluorescence imaging with IVIS. Data are expressed as the percentage of total radiant efficiency across individual organs.

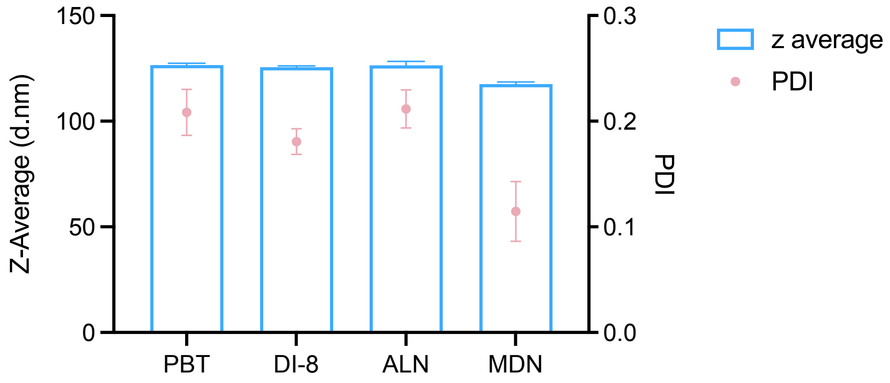


**Supplementary Figure 7. Z-average size and PDI of the four selected LNP formulations measured by DLS (n = 3) after dialysis.** Data are presented as mean ± S.D. See **Supplementary Table 2** for composition details as well as zeta potential and encapsulation efficiency of the selected LNP formulations.

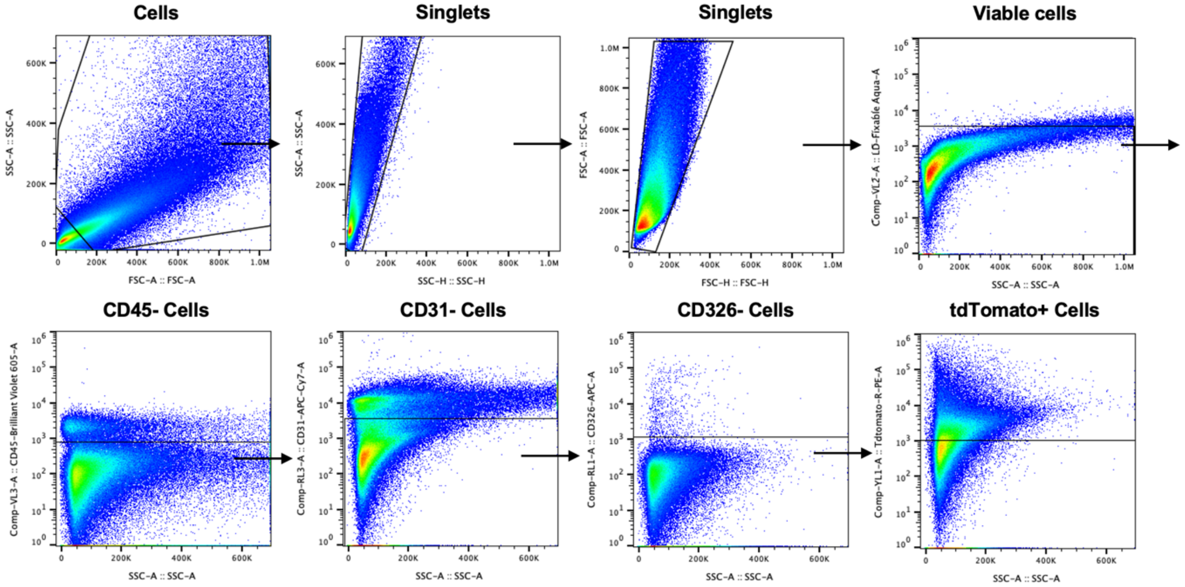


**Supplementary Figure 8. Gating strategy for flow cytometry plots for assessment of liver cell transfection by the selected LNP formulations.** Initially, cells isolated from the livers were selected using SSC-A and FSC-A parameters, followed by singlet selections with SSC-A/SSC-H and FSC-A/FSC-H plots. Viable cells were identified and selected based on the live/dead Fixable Aqua-A and SSC-A plot. Next, the immune cell (CD45), endothelial cell (CD31), and epithelial cell (CD326) populations were excluded, followed by selection of tdTom$^{+}$ hepatocytes.

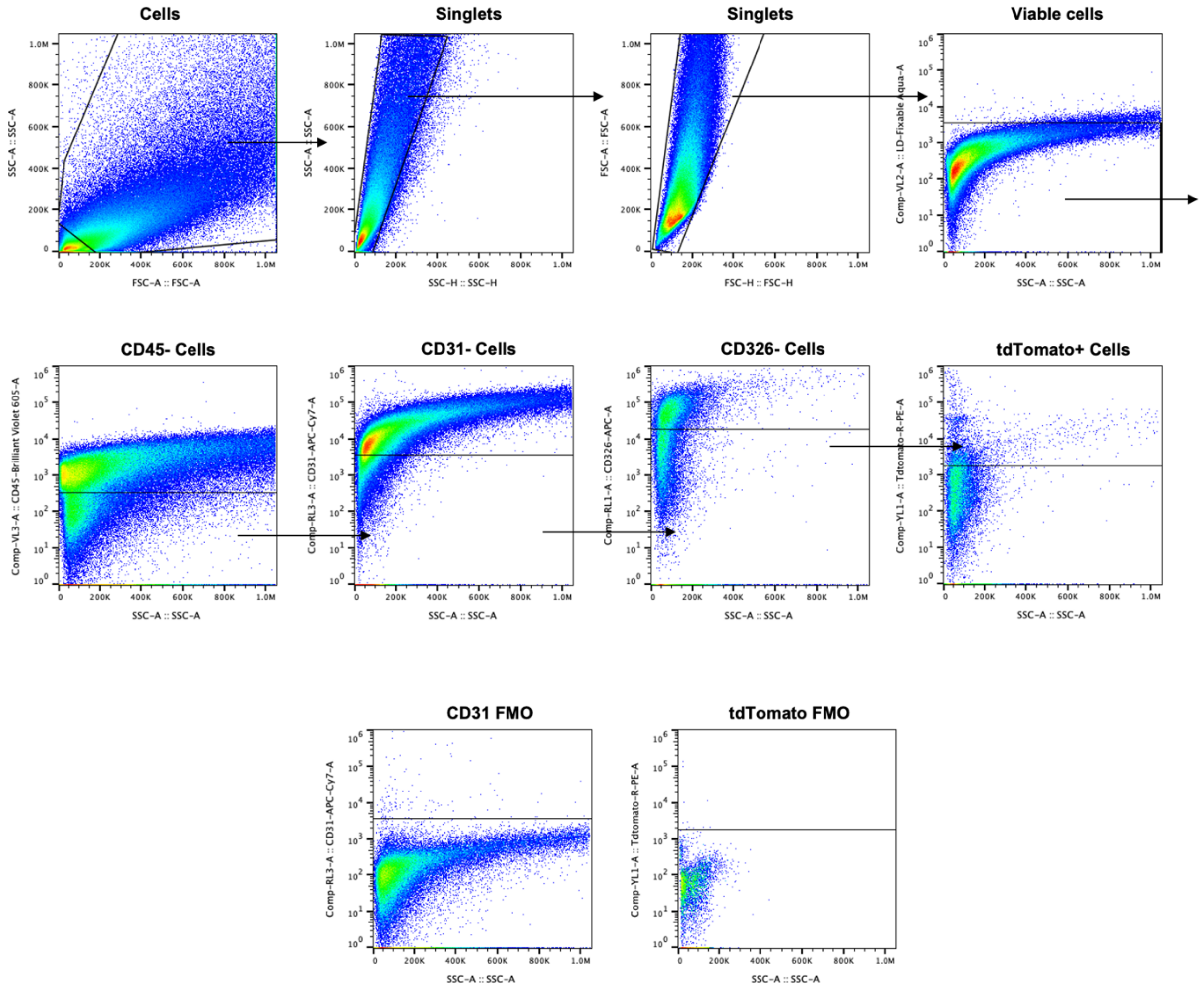


**Supplementary Figure 9. Gating strategy for flow cytometry plots for assessment of lung cell transfection by the selected LNP formulations.** Initially, cells isolated from the lungs were selected using SSC-A and FSC-A parameters, followed by singlet selections with SSC-A/SSC-H and FSC-A/FSC-H plots. Viable cells were identified and selected based on the live/dead Fixable Aqua-A and SSC-A plot. Next, the immune cell (CD45), endothelial cell (CD31), and epithelial cell (CD326) populations were excluded, followed by the selection of tdTom$^+$ cells. The CD31 FMO (Fluorescence Minus One) control for CD31 endothelial cell gating and tdTomato FMO control for tdTom$^+$ cell gating are included as references, as the population separation is not distinct.

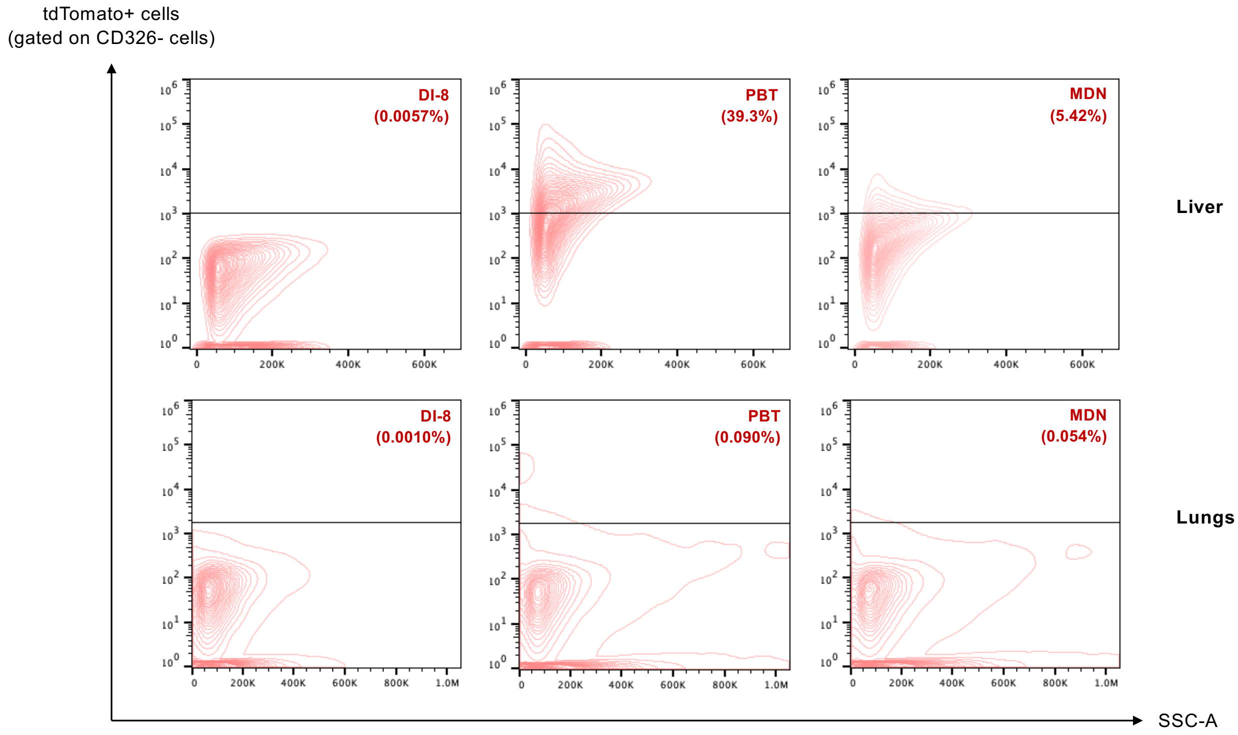


**Supplementary Figure 10. Representative flow cytometry plots for assessments of cell transfection in the liver and lungs by the selected LNP formulations.** C57BL/6 mice were given one i.m. injection of PBT, MDN, or DI-8 LNPs loaded with mCre (30 ug mCre per mouse). Successful mRNA delivery and targeting of the floxed STOP sequence turns on tdTomato expression. Mice were sacrificed on Day 7 post-injection for analysis. The livers and lungs were harvested and homogenized into cell suspension. tdTom$^{+}$ cell percentages within each organ were quantified via flow cytometry. Representative flow cytometry plots for tdTom$^{+}$ hepatocytes in the liver and tdTom$^{+}$ cells in the lungs seven days post-injection are shown. The gating strategies are shown in **Supplementary Figs. 8** and **9.**

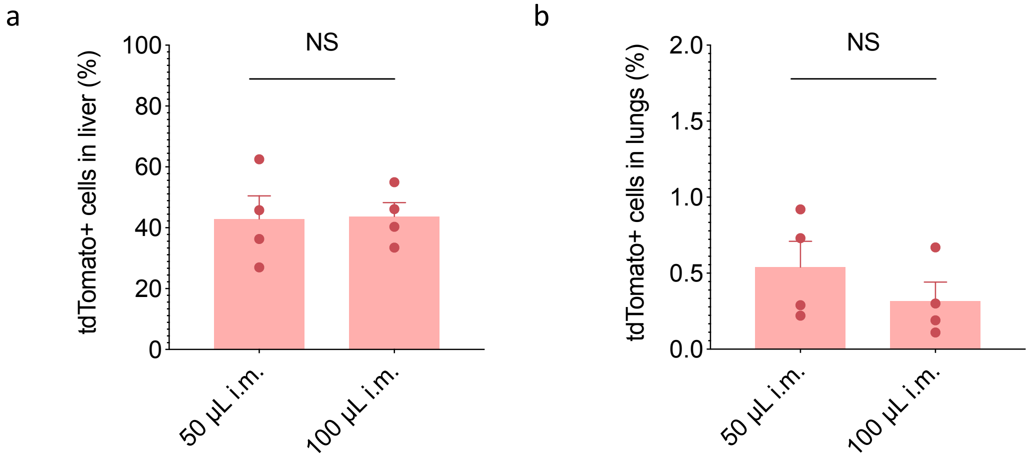


**Supplementary Figure 11. Percentage of tdTom+ cells in the liver and lungs following LNP injection by different administration routes and volumes.** C57BL/6 mice were given one i.m. injection (100 μL or 50 μL) of PBT LNPs loaded with mCre (30 μg mCre per mouse). Successful mRNA delivery and targeting of the floxed STOP sequence turns on tdTomato expression. Mice were sacrificed on Day 7 post-injection for analysis. Livers and lungs were harvested and homogenized into cell suspension. tdTom$^{+}$ cell percentages within each organ were quantified via flow cytometry. Percentages of tdTom$^{+}$ cells in the liver **(a)** and lungs **(b)** are shown. The gating strategies are shown in **Supplementary Figs. 8** and **9.**

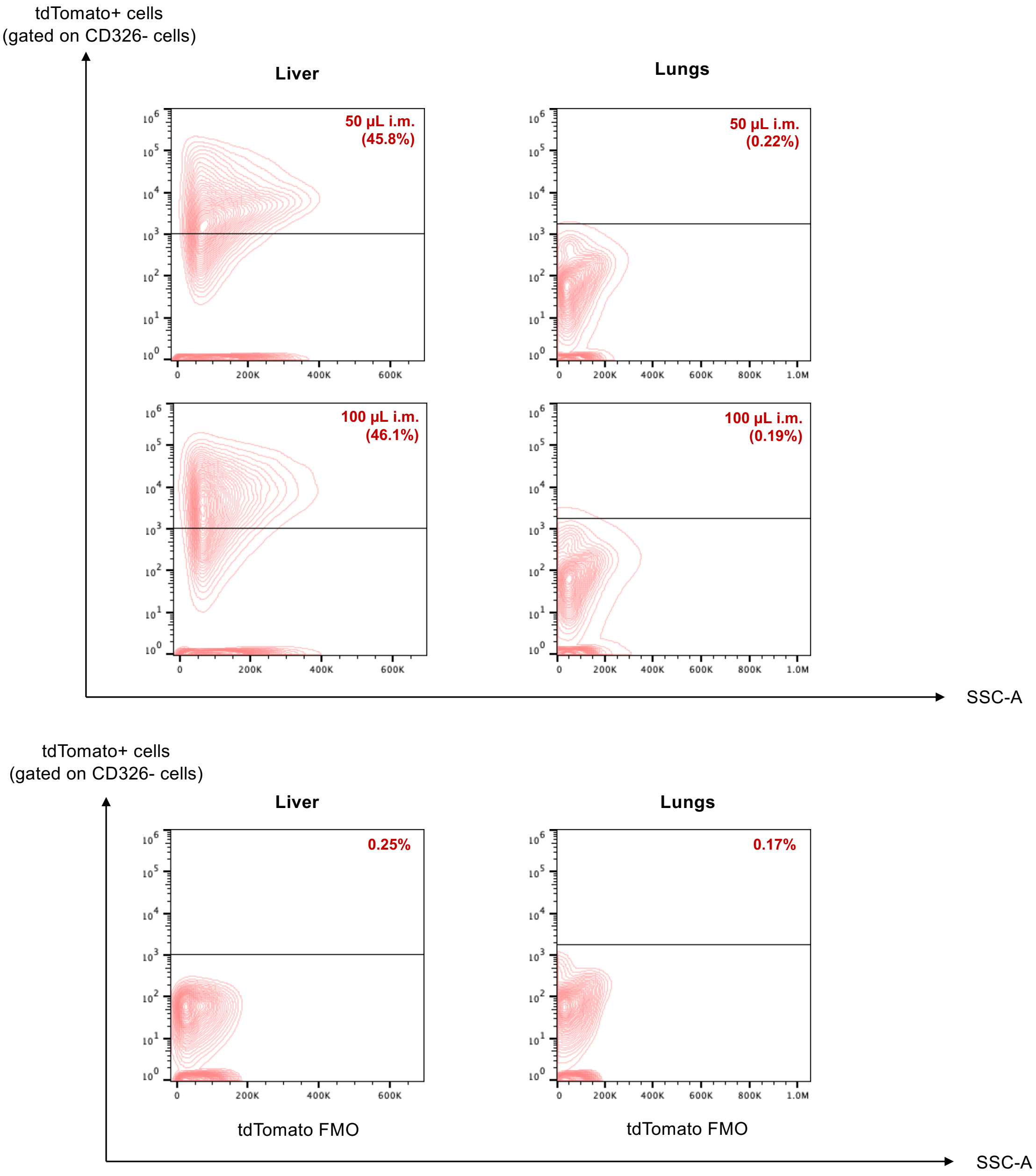


**Supplementary Figure 12. Representative flow cytometry plots for tdTom+ cells in the liver and lungs following LNP injection by different administration routes and volumes.** C57BL/6 mice were given one i.m. injection (100 μL or 50 μL) of PBT LNPs loaded with mCre (30 μg mCre per mouse). Successful mRNA delivery and targeting of the floxed STOP sequence turns on tdTomato expression. Mice were sacrificed on Day 7 post-injection for analysis. Livers and lungs were harvested and homogenized into cell suspension. $tdTom^+$ cell percentages within each organ were quantified via flow cytometry. Percentages of $tdTom^+$ cells in the liver and lungs are shown. The tdTomato FMO (Fluorescence Minus One) controls for $tdTom^+$ transfected cell gating for hepatocytes and lung cells are included as references, as the population separation is not distinct. The gating strategies are shown in **Supplementary Figs. 8** and **9.**

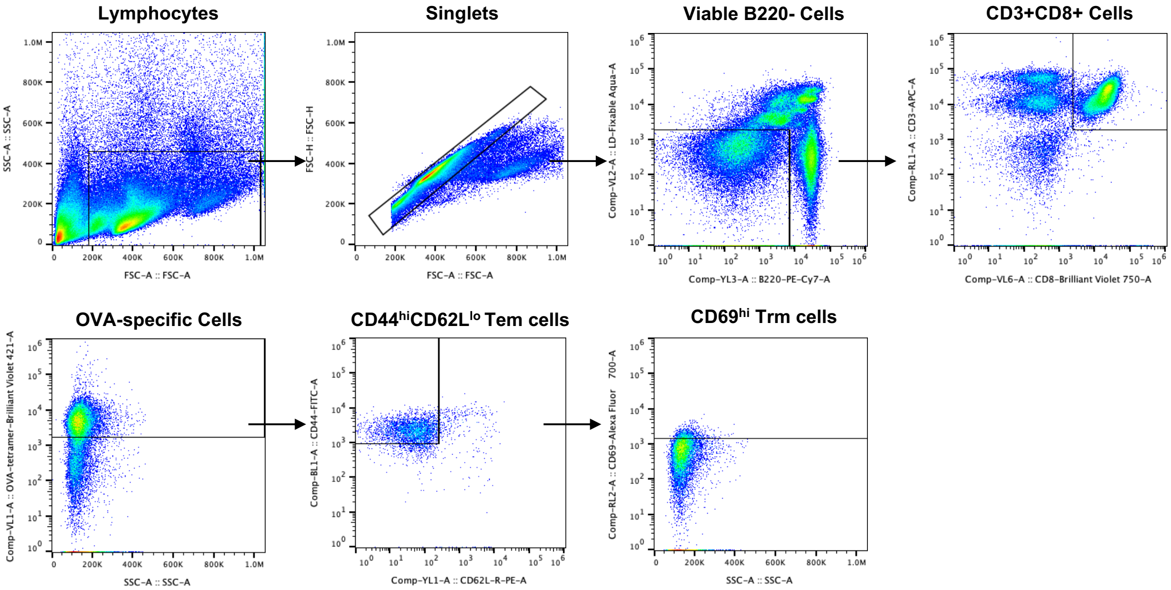


**Supplementary Figure 13. Gating strategy for flow cytometry plots for T cell responses after vaccination.** Initially, lymphocytes isolated from the liver, lungs, and spleen were selected using SSC-A and FSC-A parameters, followed singlet selection with FSC-H and FSC-A plot. Viable cells were identified and selected based on the live/dead Fixable Aqua-A staining, and $B220^{+}$ B cell population was excluded from downstream analysis. Next, $CD3^{+}$ and $CD8^{+}$ cell population was selected for antigen-specific T cells, in which $CD44^{hi}CD62L^{lo}$ Tem-like cells were identified, followed by the subsequent $CD69^{hi}$ Trm-like cell identification.

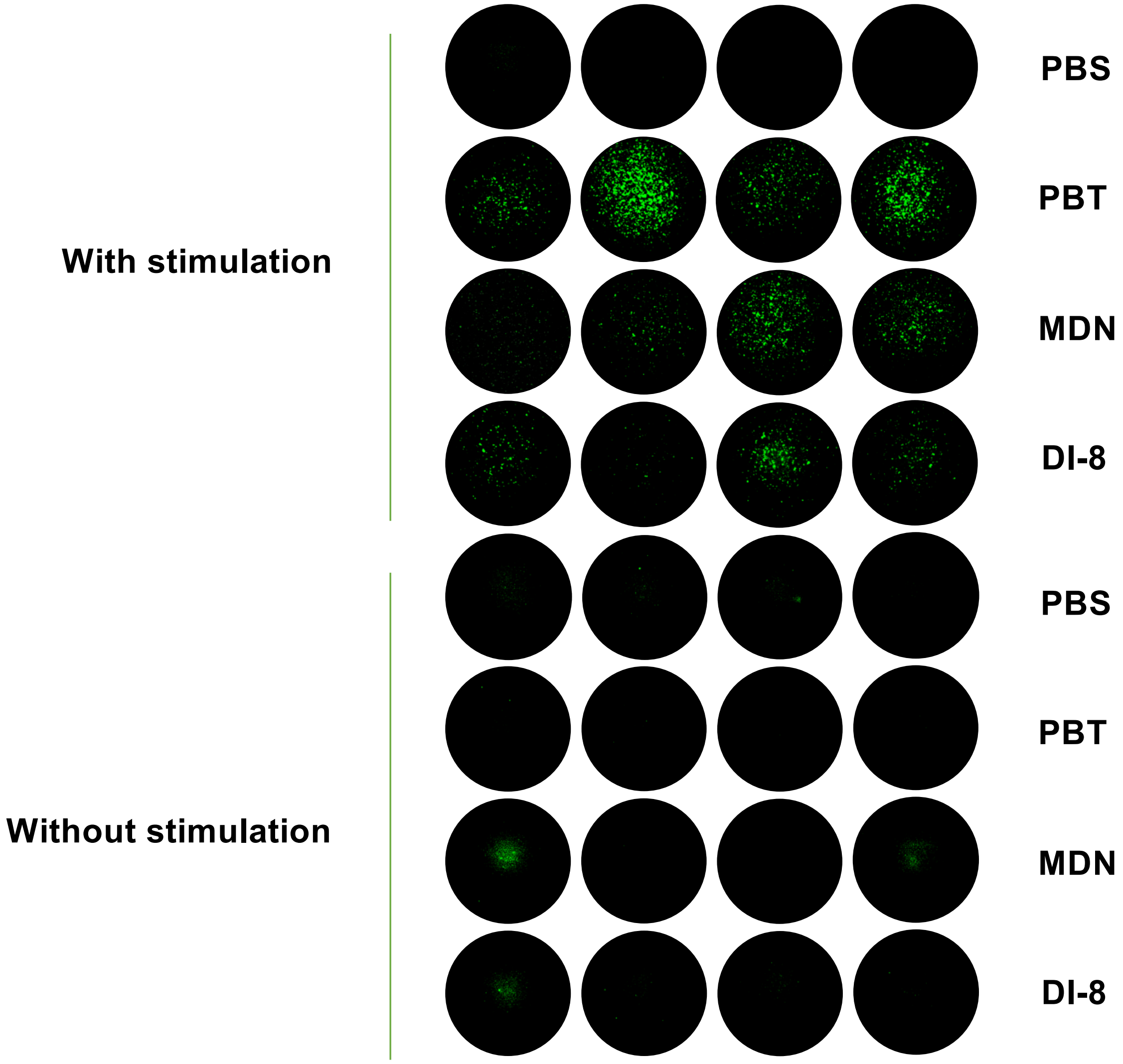


**Supplementary Figure 14. Representative images of IFN-γ-secreting cells in the liver at the short-term time point (Day 28) from the FluoroSpot assay.** Frequency of IFN-γ-secreting cells among restimulated lymphocytes assessed via FluoroSpot. Lymphocytes were restimulated *in vitro* with SIINFEKL peptide (2 μg/mL SIINFEKL).

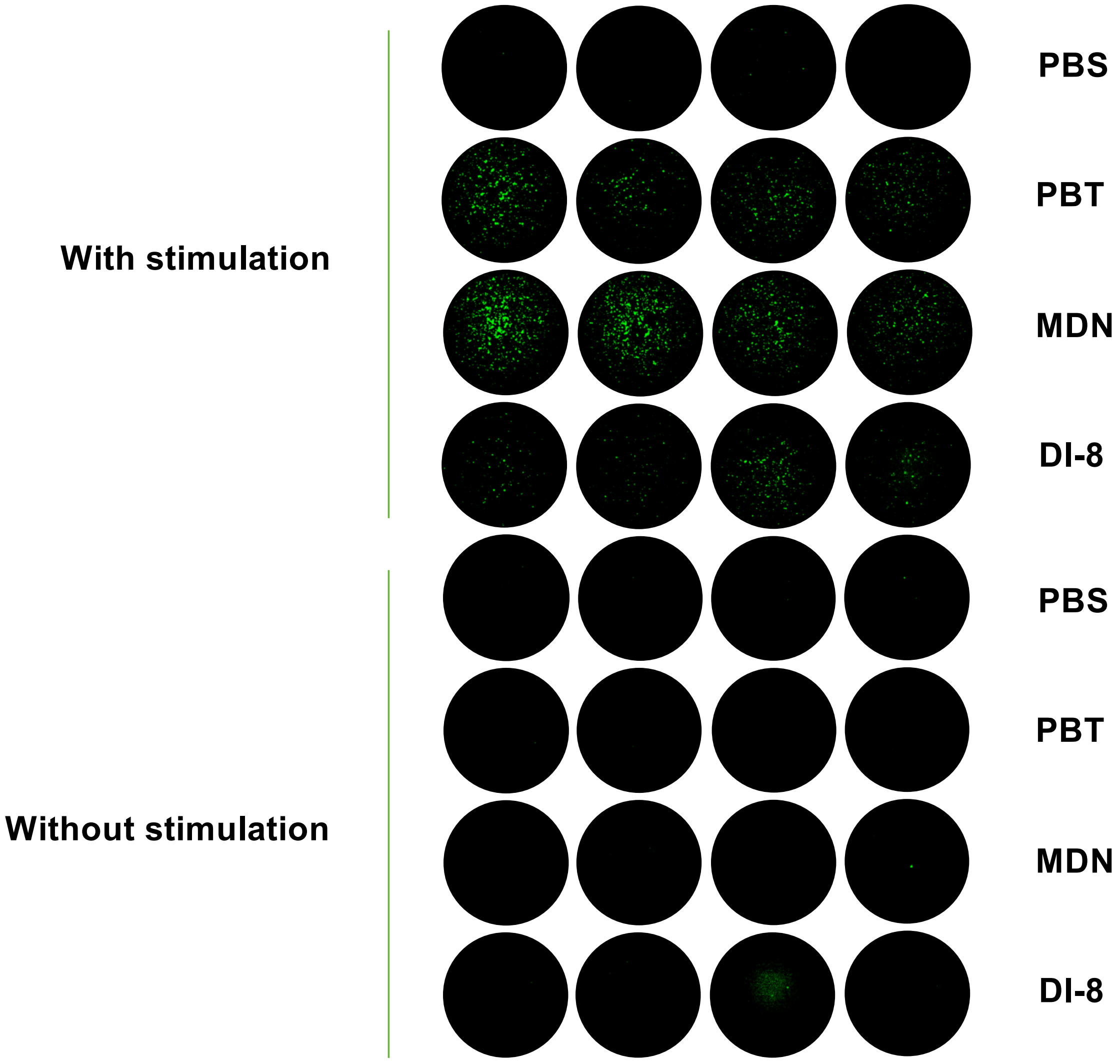


**Supplementary Figure 15. Representative images of IFN-γ-secreting cells in the lungs at the short-term time point (Day 28) from the FluoroSpot assay.** Frequency of IFN-γ-secreting cells among restimulated lymphocytes assessed via FluoroSpot. Lymphocytes were restimulated *in vitro* with SIINFEKL peptide (2 μg/mL SIINFEKL).

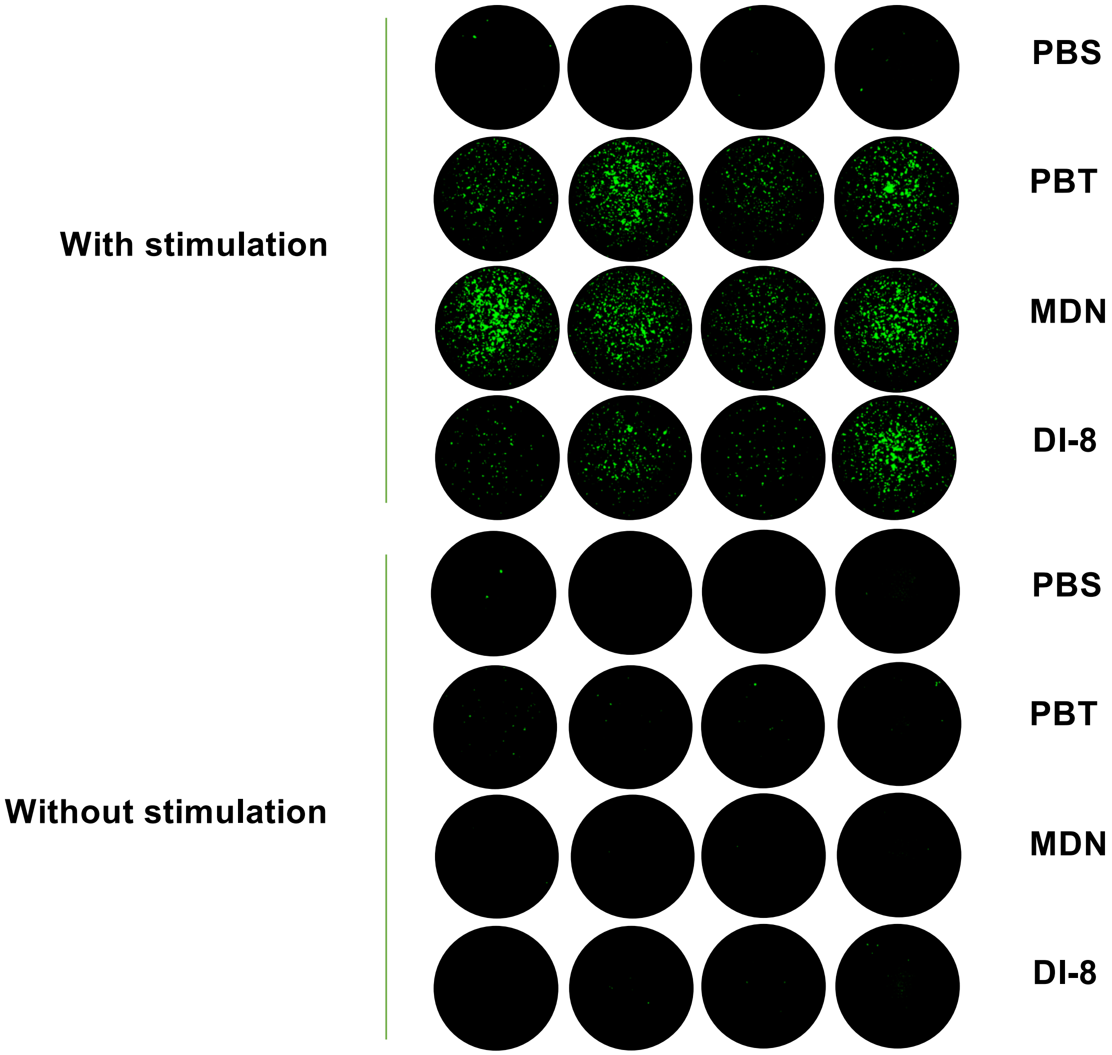


**Supplementary Figure 16. Representative images of IFN-γ-secreting cells in the spleen at the short-term time point (Day 28) from the FluoroSpot assay.** Frequency of IFN-γ-secreting cells among restimulated lymphocytes assessed via FluoroSpot. Lymphocytes were restimulated *in vitro* with SIINFEKL peptide (2 μg/mL SIINFEKL).

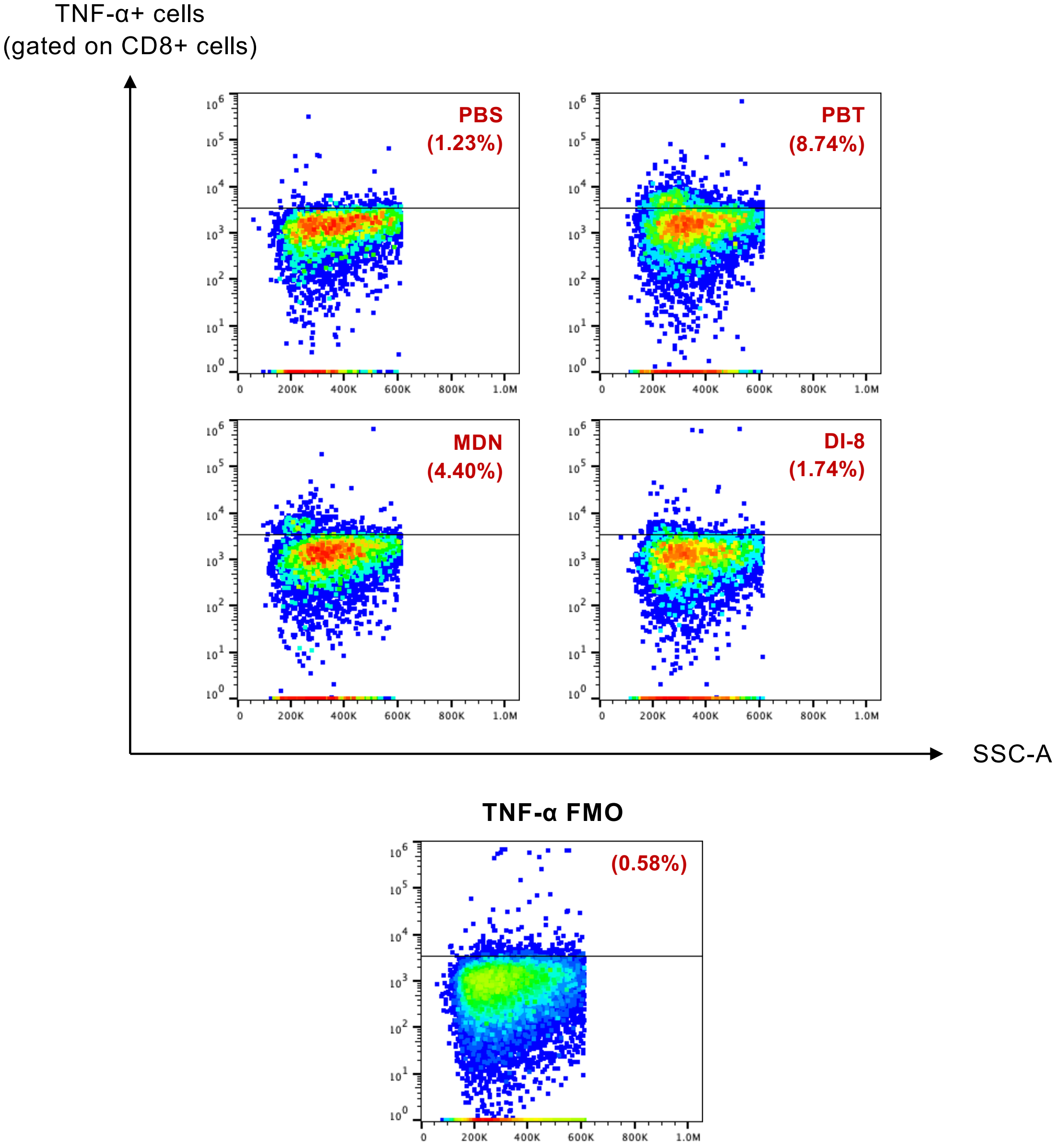


**Supplementary Figure 17. Representative flow cytometry plots for analysis of CD3+CD8+TNFα+ cells in the spleen at the long-term time point (Day 90).** Lymphocytes isolated from the spleen were restimulated *in vitro* with OVA and SIINFEKL peptide (100 μg/mL OVA and 2 μg/mL SIINFEKL) for 12 h and assessed via intracellular cytokine staining and flow cytometry and to determine the percentages of $CD3^{+}CD8^{+}TNF\text{-}\alpha^{+}$ cells.

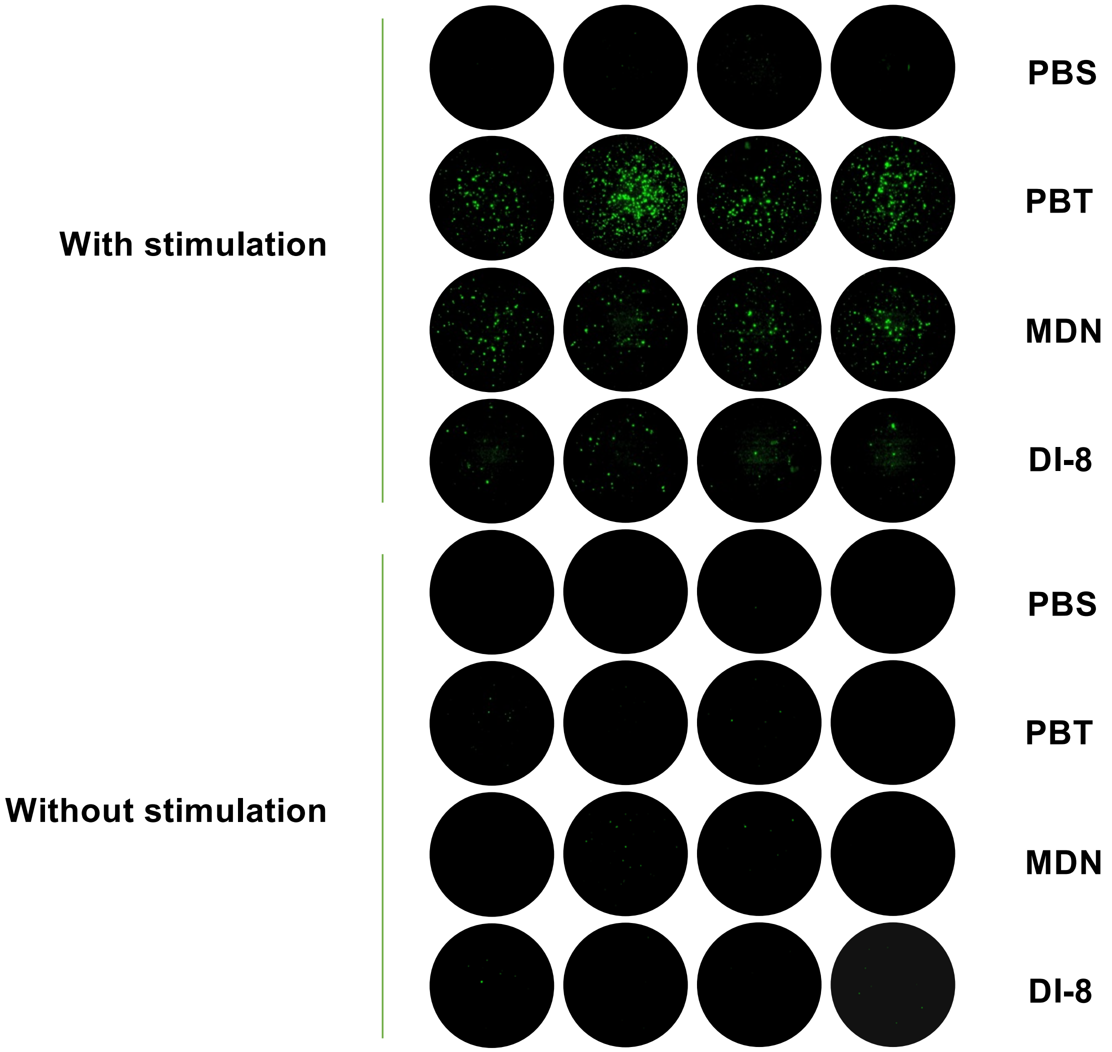


**Supplementary Figure 18. Representative images of IFN-γ-secreting cells in the spleen at the long-term time point (Day 90) from the FluoroSpot assay.** Frequency of IFN-γ-secreting cells among restimulated lymphocytes assessed via FluoroSpot. lymphocytes were restimulated *in vitro* with SIINFEKL peptide (2 μg/mL SIINFEKL).

**Supplementary Table 1. Anti-mouse antibodies used in flow cytometry panels. Marker, fluorophore, catalogue number, source, and concentration are indicated.**

| Antigen | Fluorophore | Catalogue # | Source | Concentration |
|---|---|---|---|---|
| CD8a | APC-Cy7 | 100714 | Biolegend | 1:200 dilution |
| OVA SIIGFEKL (Tetramer) | Brilliant Violet 421 | N/A | NIH tetramer core facility | 1:400 dilution |
| CD3 | APC | 100236 | Biolegend | 1:100 dilution |
| Live/Dead | Live/Dead Fix Aqua | L34957 | Thermo Fisher | 1:1000 dilution |
| CD44 | FITC | 103006 | Biolegend | 1:100 dilution |
| CD62L | PE | 161204 | Biolegend | 1:100 dilution |
| CD45 | FITC | 103108 | Biolegend | 1:250 dilution |
| CD31 | APC-Cy7 | 102440 | Biolegend | 1:100 dilution |
| CD326 | Brilliant Violet 605 | 118227 | Biolegend | 1:100 dilution |
| CD45 | Brilliant Violet 421 | 103134 | Biolegend | 1:250 dilution |
| CD3 | PE | 100206 | Biolegend | 1:200 dilution |
| CD8 | FITC | 100706 | Biolegend | 1:200 dilution |
| Fixable Live/Dead | Live/Dead Fix Near IR (780) | L10119 | Thermo Fisher | 1:1000 dilution |
| CD8a | Brilliant Violet 750 | 747134 | Biolegend | 1:200 dilution |
| CD45R | PE-Cy7 | 103222 | Biolegend | 1:100 dilution |
| CD69 | Alexa Fluor 700 | 104539 | Biolegend | 1:50 dilution |
| TNF alpha | Alexa Fluor 700 | 506338 | Biolegend | 1:100 dilution |
| CD3 | Brilliant Violet 421 | 100228 | Biolegend | 1:200 dilution |
| CD326 | APC | 118214 | Biolegend | 1:100 dilution |

**Supplementary Table 2. Composition details and characterization of the five evaluated LNP formulations.**

| Formulation Code | PBT | C10 | DI-8 | ALN | MDN |
|---|---|---|---|---|---|
| *Composition (molar ratio):* | | | | | |
| Ionizable lipid | 46.3 | 40 | 36.36 | 49.74 | 50 |
| Helper lipid | 9.4 | 40 | 3.64 | 10.26 | 10 |
| Cholesterol | 42.7 | 19.96 | 59.88 | 38.46 | 38.5 |
| DMG-PEG2000 | 1.6 | 0.04 | 0.12 | 1.54 | 1.5 |
| N/P ratio | 6 | 4 | 8 | 4 | 6 |
| *Formulation features:* | | | | | |
| Z-average diameter (nm) | 126.4 ± 0.6 | 162.9 ± 2.6 | 125.6 ± 0.6 | 126.6 ± 0.9 | 117.6 ± 1.0 |
| Average PDI | 0.212 ± 0.018 | 0.209 ± 0.009 | 0.181 ± 0.012 | 0.208 ± 0.022 | 0.115 ± 0.028 |
| Average Zeta potential (mV) | -2.92 ± 0.20 | -11.0 ± 2.30 | -1.25 ± 0.50 | -2.95 ± 0.51 | -2.80 ± 0.71 |
| Average EE% | 96.45 | 99.03 | 99.96 | 99.95 | 99.85 |
